# Collective directional memory controls the range of epithelial cell migration

**DOI:** 10.1101/2023.12.08.570811

**Authors:** Helena Canever, Hugo Lachuer, Quentin Delaunay, François Sipieter, Nicolas Audugé, Philippe P. Girard, Nicolas Borghi

**Author notes:** These authors contributed equally (Helena Canever, Hugo Lachuer).

## Abstract

Cell migration is a fundamental behavior in multicellular development, regeneration, and homeostasis, which is deregulated in cancer. Epithelial cells migrate individually when isolated and collectively within a tissue. However, how interactions between cells affect their ability to explore space and their sensitivity to guidance signals is poorly understood. We show that isolated cells that are persistent random walkers adopt a super-diffusive behavior in an epithelium. The effect is stronger than external guidance cues and enables cells to reach greater distances than when isolated. This behavior is consistent with a fractional Brownian motion that emerges from velocity coordination between neighboring cells with intact intercellular adhesion. Furthermore, we show how the molecular stability and mechanosensitivity of adhesion complexes, both linked to the ability of the adhesion protein vinculin to dimerize, ultimately regulate the speed of collective migration and the sensitivity to guidance signals. Together, our results show how cell speed, persistence, and directionality define the efficiency of spatial cell exploration on short, intermediate, and long time scales, respectively.

## Introduction

In multicellular organisms, cell migration is an essential process in tissue morphogenesis, homeostasis, and repair. Deregulation of cell migration is also involved in the metastatic invasion of cancers (1). Cell migration occurs in different ways, depending on the intrinsic properties of cells and how they interact with each other and their environment. As a result, cells may migrate individually or collectively, guided by local or global signals (2).

For these reasons, how cells explore their environment and how cell interactions and guidance cues affect this exploration are fundamental questions that have motivated abundant research for over 70 years (3, 4). Since then, various models have proven useful in describing the exploration of space by individually migrating cells. Directional persistence has emerged as a consistent, though highly variable, feature arising from the non-Gaussian distribution of cell displacements, temporal velocity correlations, or both , and manifests itself through a finite persistence time or anomalous diffusion depending on the type of cell or the context (5–8). When migrating collectively, neighbor cells typically exhibit correlated velocities that depend on their interactions (9–11). This coordination and the diverse mechanisms underlying it have been the subject of intense scrutiny for the past decade (12). In comparison, how each cell in the collective explores its environment has received much less attention, with a few exceptions (13–15). Remarkably, it remains unclear whether neighbor interactions actually help or hinder the exploratory capacity or sensitivity to guidance cues of each cell in the collective, much less whether and how they quantitatively or qualitatively affect the underlying cellular and molecular mechanisms of migration.

Adherent cells form Focal Adhesions (FAs) on their substrate, which they continuously assemble at their front and disassemble at their back during migration (16). There, mechanosensory proteins, such as vinculin, provide force transmission from the contractile cytoskeleton (17). In cohesive tissues, cells also adhere to each other through cadherin-based Adherens Junctions (AJs) where *α*-catenin and vinculin enable dynamic connections with the cytoskeleton and force transmission (18). Vinculin comprises a head and a tail domain whose intramolecular interaction competes with intermolecular dimerization and binding to the actin cytoskeleton of its tail, and with binding of its head to adapter proteins in FAs and AJs (19). In AJs, *α*-catenin is this adapter, which also binds actin by its own tail (20). Vinculin deficiencies impact the migration of cells as individuals, in a collective, and with guidance cues, although in a cell type- and mutation-dependent manner (11, 21–32). Alpha-catenin deficiencies have an impact on guided and coordinated collective migration, involving the interaction of *α*-catenin with vinculin in AJs (11, 26, 33, 34). However, the lack of comparable metrics, scales and molecular, cellular and migratory contexts between studies makes it difficult for a general picture to emerge, where the functions of proteins in regulating the molecular stability and mechanosensitivity of adhesions could explain how they control the intrinsic speed of cells, their persistence, and their responses to interactions with neighbors and guidance cues.

Here, we address this issue by comparing the exploratory ability of epithelial cells as individuals, in the collective and exposed to a guidance cue, while genetically disrupting the interactions of cells with their substrate and neighbors and monitoring the associated effects on the dynamics and mechanics of adhesion complexes. We show that individual epithelial cells migrate with transient directional memory and that vinculin regulates their speed by synergizing FA molecular stability and cytoskeletal tension transmission through its dimerization and actin-binding abilities, respectively. In contrast, cells that migrate collectively acquire long-term directional memory, which specifically requires *α*-catenin in AJs and results from coordination of velocities between neighboring cells. For this reason, we define it as a collective directional memory. In this context, vinculin additionally controls the speed of cells from their AJs through its impact on their mobility, but in a manner specifically linked to its ability to dimerize. Overall, this results in a spatial exploration that is less efficient than that of individual cells on time scales less than ≈3hrs but more efficient beyond that. Finally, cell guidance by an epithelial wound also requires *α*-catenin, but the directional bias is too small to significantly affect the efficiency of spatial exploration for hours. In this context of guided collective cell migration, the balance in mechnosensitivity between FAs and AJs, which is also linked to the ability of vinculin to dimerize specifically in AJs, regulates wound healing efficiency.

## Results

### Single epithelial cells are persistent random walkers

To characterize the motion of single epithelial cells, we tracked nucleus-stained MDCK cells overnight, sparsely plated on collagen-coated glass coverslips, which offer a reasonable approximation of the basement membrane(35–37) (see Materials and Methods)(Fig. 1A, movie 1). Based on the Akaike information criterion, we found that the average Mean Square Displacement as function of lag time Δ*t* (MSD, see Materials and Methods) was better described by a persistent random walk (38, 39) than anomalous diffusion: the MSD transitioned from ballistic (MSD ∼ Δ*t*^2^, Δ*t* → 0) to pure diffusion (MSD ∼ Δ*t*, Δ*t* → ∞) with a persistence time *P* of the order of 10min, rather than scaling as Δ*t*^*α*^ with *α* ≠ 1 (Fig. 1B; Table S1). The ability to extract a finite persistence time from such an ensemble and time-averaged measurement supports the view that there is no substantial heterogeneity in persistence time among the cell population and along cell trajectories. To confirm the underlying walk model, we examined the step length *R* for the step durations *τ* = 10min and *τ* = 3hrs. In both cases, *R* followed a Rayleigh distribution, in agreement with a Gaussian walk regardless of step duration (Fig. 1C, S1A). Moreover, despite temporal variations in the step length distributions, no aging was observed. Finally, we examined the angular memory, which we define as a modified angular correlation in which we consider the angular deviation *θ* of each elementary step of duration 10min from its immediate previous step of duration *τ* = Δ*t* (see Materials and Methods). We found that the distribution of *θ* across the cell population was skewed towards small angles at small Δ*t* and uniform at large Δ*t*, consistent with a persistent random walk in which the angular correlation vanishes beyond *P* (Fig. S1B). Moreover, the lack of an inflection point for any Δ*t* indicated no substantial heterogeneity in the cell population. Consistently, the angular memory decreased to 0 with increasing Δ*t* (Fig. 1D). Overall, these results show that single epithelial cells migrate as persistent random walkers, displaying transient directional persistence overcome by pure diffusion on longer time scales.

**Fig 1.**
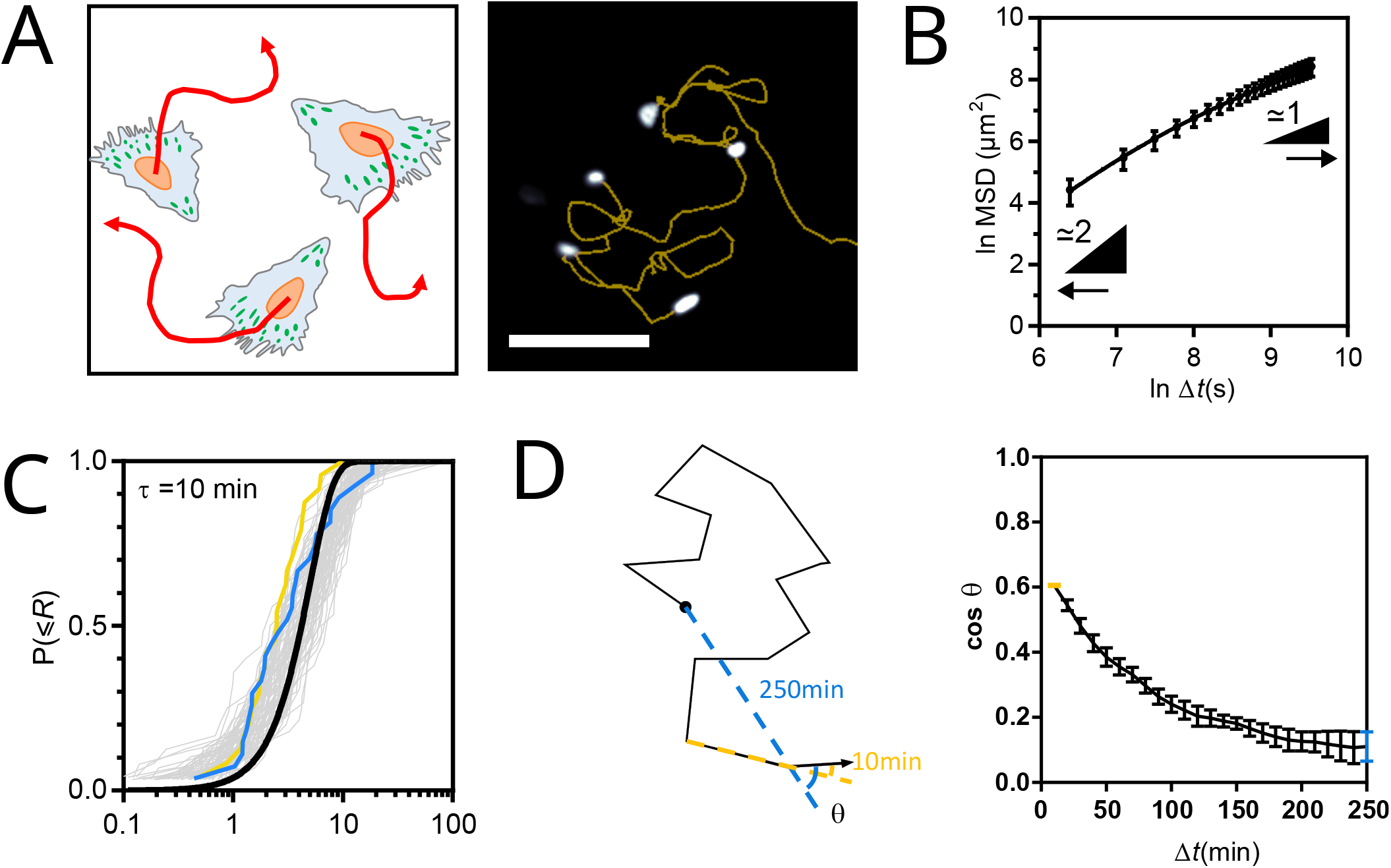
(A) Left: Schematic representation of the single cell migration assay. Right: fluorescence image of nucleus-stained cells and their migration tracks in yellow. Scale bar = 100*µ*m. (B) MSD(Δ*t*) of WT cells fit with the Fürth model of a persistent random walk, the better model according to the Akaike information criterion (see Materials and Methods). (C) Cumulative distributions of the step length *R* for *τ* = 10min. One grey curve per frame pair along a movie. The yellow and blue curves correspond to the first and last frames, respectively. The black line is a global fit of all curves with a cumulative Rayleigh distribution function. (D) Left: Schematic migration track and angular deviation *θ*(Δ*t*) (see Materials and Methods) at Δ*t* = 10min (yellow) and Δ*t* = 250min (blue). Right: Angular memory (cos *θ*) between Δ*t* = 10min (yellow dot) and Δ*t* = 250min (blue dot). The black line connects neighboring data points. Data of (B-D) from 94 cell tracks of 2 independent experiments. Mean *±* SEM.

To investigate the robustness of this behavior, we examined the effects of overexpressing full-length (FL) vinculin or vinculin mutants, also equipped with a tension sensor module (40, 41). Specifically, we considered the T12 mutant with charge-to-alanine mutations in the tail that impair the head-tail interaction (42), the Y1065E (YE) mutant with a phsophomimetic mutation in the tail that impairs the tail-tail interaction (43) and the Tailless mutant (TL) lacking amino acids beyond 883 that are required for all tail interactions (44) (Fig. S2A). All constructs localized to FAs in stable cell lines (Fig. S2B,C), which exhibited an MSD better described by a persistent random walk model than by a pure or superdiffusion model (Fig. 2A, Table S1). The effects of mutant expression on persistence times and diffusion coefficients were variable and did not show a correlation (Fig. S2D). In addition, the diffusion coefficient *D* scaled well with the squared apparent speed *v*_s_ with a common persistence time for all cell lines (Fig. S2E), showing that cell lines differed primarily in apparent speed rather than persistence. Specifically, TL and YE mutants with impaired actin-tail or tail-tail interactions decreased cell speed, while the T12 mutant with altered head-tail interaction increased cell speed relative to FL vinculin (Fig. 2B). This could not be explained by differences in expression levels between cell lines since the effects of YE and TL were stronger than those of FL despite an equal or lower expression, and those of T12 completely opposite despite a higher expression (Fig. S2F). Rather, this indicated a dominant effect, consistent with previous reports for T12, TL and other mutants (44). Overall, these results show that the persistence, but not the speed of single migrating cells is robust to mutations in the FA protein vinculin.

**Fig 2.**
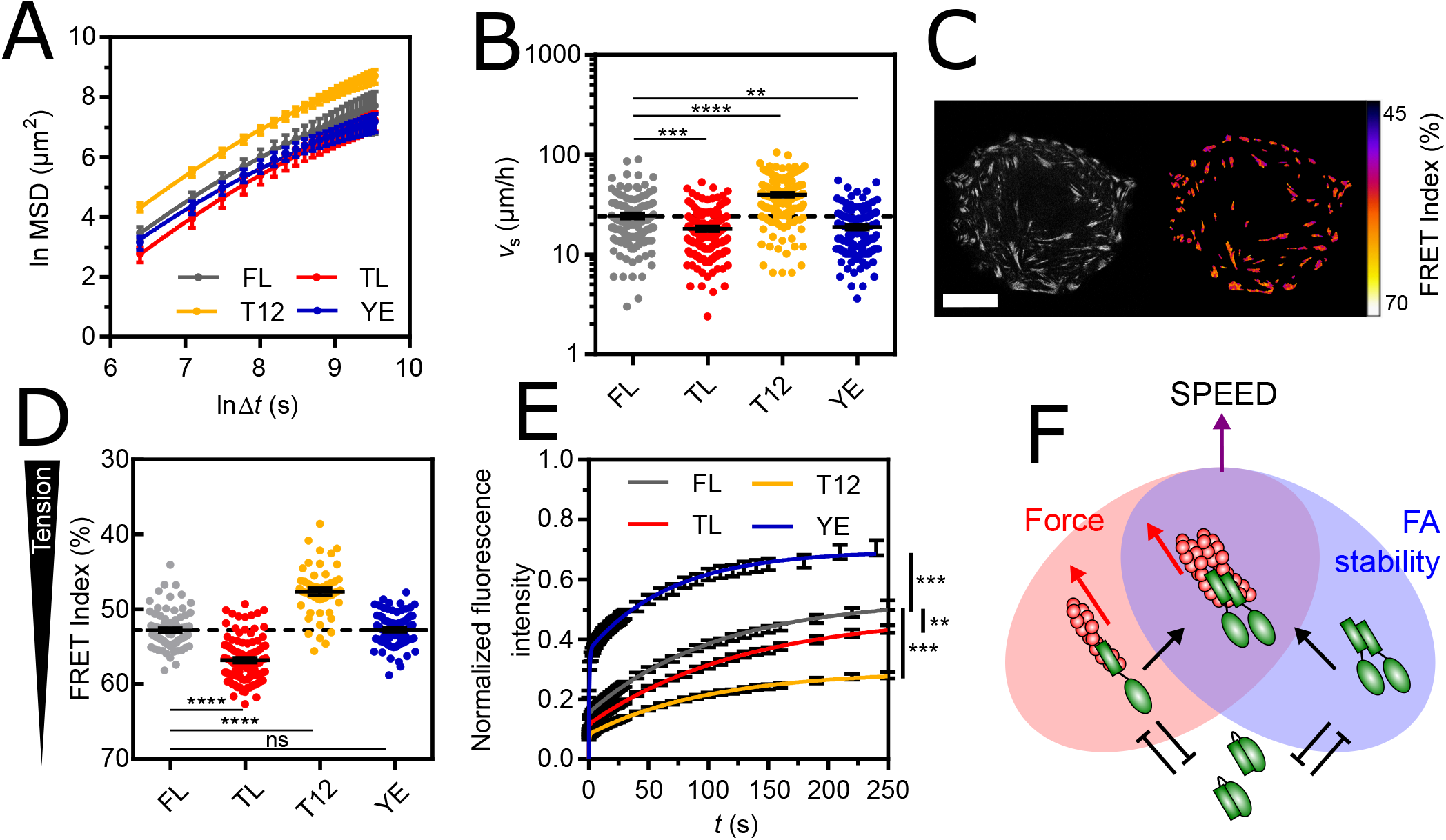
(A) MSD(Δ*t*) of vinculin FL and mutant cells fit with the Fürth model of persistent random walk. (B) Apparent speed of single, vinculin FL and mutant cells. The dashed line indicates the mean of FL as a reference. (C) Left: direct fluorescence of vinculin TSMod in a single cell. Right: FRET Index map (see Materials and Methods). Scale bar = 20 *µ*m. (D) FRET Index of TSMod in the FL, TL, T12 and YE constructs in FAs. The dashed line indicates the mean of FL as a reference. (E) Fluorescence recovery after photobleaching of FL, TL, T12 and YE constructs in FAs fit with a 2-phases association model. (F) Proposed Boolean model as a Venn diagram of the possible conformations and associated interactions of intact vinculin, and their individual and combined consequences on adhesion and migration: at FAs, the head-tail interaction of vinculin (in green) competes with both tail dimerization and actin filaments (in red) binding, which provide FA molecular stability (blue ellipse, from panel E) and force transmission (red ellipse, from panel D), respectively, together promoting cell speed (ellipses overlap, from panel B). Data of (A,B) from 125 (FL), 129 (TL), 193 (T12) and 110 (YE) cell tracks of 3 independent experiments. Data of (D) from 81 (FL), 102 (TL), 52 (T12) and 84 (YE) cells of 4 (FL) and 3 (TL, T12, YE) independent experiments. Data of (E) from 105 (FL), 112 (TL), 118 (T12) and 65 (YE) cells of 2 independent experiments. Mean *±* SEM. Kruskal-Wallis and Dunn’s tests between each mutant and FL (B,D). Extra-sum-of-squares F test with shared immobile fraction between each mutant and FL as the null hypothesis (E). Benjamini-Krieger-Yekuteli-corrected p-values. *: p<0.05, **: p<0.01, ***: p<0.001, ****: p<0.0001.

### The combination of Focal Adhesion molecular stability and force transmission from the cytoskeleton predicts the migration speed of single cells

The transmission of cytoskeletal forces through FAs is essentially based on vinculin (44–46). Thus, to identify the underlying cellular mechanisms and molecular determinants of speed regulation by FAs, we took advantage of the tension sensor module in vinculin and its mutants to assess their mechanical state by Molecular Tension Microscopy (Fig. 2C) (see Materials and Methods). By design, a decrease in FRET index reports an increase in tension per molecule (40). We found that alteration of actin-tail and head-tail interactions (TL and T12) decreased and increased vinculin tension relative to FL vinculin, respectively (Fig. 2D), as seen in other cell types (40, 41). In contrast, alteration of the tail-tail interaction (YE) did not substantially affect vinculin tension, meaning that the ability of vinculin to dimerize is unnecessary for force transmission (Fig. 2D). Next, we investigated a potential relationship between vinculin tension and single cell speed across vinculin constructs, but found no significant correlation (Fig. S2G), indicating that single cell speed is not simply a result of vinculin force transmission.

The stability of the core cytoskeletal linkers of FAs is based on vinculin (30, 47). Therefore, we investigated the effects of mutations on vinculin turnover by Fluorescence Recovery After Photobleaching (FRAP, Fig. S2H, movie 2) (see Materials and Methods). We found that TL and T12 mutants, both of which impair head-tail interaction, had a significantly higher immobile fraction than FL vinculin (Fig. 2E), consistent with previous studies in other cell types (30, 44, 47–50). This shows that impaired head-tail interaction, whether associated with impaired actin-binding (TL) or increased tension (T12), is sufficient for vinculin stability in FAs. In contrast, the YE mutant that impairs the tail-tail interaction showed a significantly smaller immobile fraction (Fig. 2E). This supports that the ability of vinculin to dimerize promotes its stability. Additionally, this shows that cell migration speed does not simply result from vinculin stability since T12 and TL induce opposite effects on migration speed while both stabilize vinculin, and T12 and YE have antagonist effects on vinculin stability while both promote cell speed.

Together, this supports that migration speed decreases upon loss of force transmission or the molecular stability of FAs (TL and YE) and increases upon strengthening of both (T12). We therefore propose that single cell migration speed results from the combination of FA molecular stability and force transmission, the former provided by the ability of vinculin to dimerize and the latter by vinculin-actin binding, and both competed for by its head-tail interaction. We provide a simple formalization with a Boolean model depicted by a Venn diagram in Fig.2F. To quantify this, we examined how 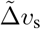, the change in cell speed across vinculin constructs re-scaled between its extreme values, depended on the corresponding changes in vinculin tension 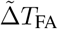 and vinculin stability 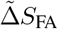, obtained from similarly re-scaled FRET index and immobile fraction measurements, respectively (see Material and Methods). We found that the optimal linear combination of 2*/*3 change in tension and 1*/*3 change in stability explained most of the change in cell speed across vinculin constructs (Fig. S2I, see Materials and Methods). Cell speed is therefore more susceptible to transmission of force in FAs than to their molecular stability.

### Collectively migrating epithelial cells are super-diffusive, fractional Brownian walkers

To assess how intercellular adhesion affects the persistent random walk of cells, we tracked MDCK cells plated at confluence on collagen-coated glass coverslips (see Materials and Methods) (Fig. 3A, movie 3). We measured the cell shape index *p* (see Materials and Methods), which quantifies departure from a circular shape of the apical cell surface and decreases when cortical tension increases in vertex models of epithelia (51). In these conditions, *p* ≈ 4.2 was well above the rigidity threshold (*q* ≈ 3.8, Fig. S3A) , which supports that the tissue behaved as a fluid with negligible resistance arising from cortical tension against cell rearrangements (52). We found that within the same time frame as in single cell migration experiments, cells no longer migrated as persistent random walkers but adopted super-diffusive behavior with an anomalous exponent of approximately 1.7 (Fig. 3B, Table S2). Remarkably, this superdiffusive behavior allowed cells to eventually reach a greater distance than single cells in the same amount of time, even though confluence also lowered their apparent speed (Fig. S3B)

To distinguish between the various walk models that could produce such a super-diffusive behavior (Lévy walk, fractional Brownian motion, or their combinations (7, 53)), we examined the step length distributions and found that they were consistent with a Gaussian walk on any time scale up to *τ* = 3hrs and showed no sign of aging (Fig. 3C, S3C). However, we found that the angular deviation *θ* of any 10min-step from the direction of the previous step was skewed toward small angles as in single cells but remained so for durations of the previous step up to *τ* = Δ*t* = 250min (Fig. S3D). This is consistent with a scale-invariant angular deviation showing no sign of heterogeneity among the cell population. On average, the angular memory never vanished to 0 within our experimental time frame, unlike that of single cells, consistent with a long-term directional memory (Fig. 3D compared to Fig. 1D). Gaussianity and long-term directional memory are characteristics of a pure fractional Brownian motion and thus rule out Lévy walks, with anomalous step distributions, and more complex scenarios involving their combination or heterogeneous behaviors in time or among the population. Cell super-diffusion thus emerges from a long-term directional memory consistent with a fractional Brownian motion.

**Fig 3.**
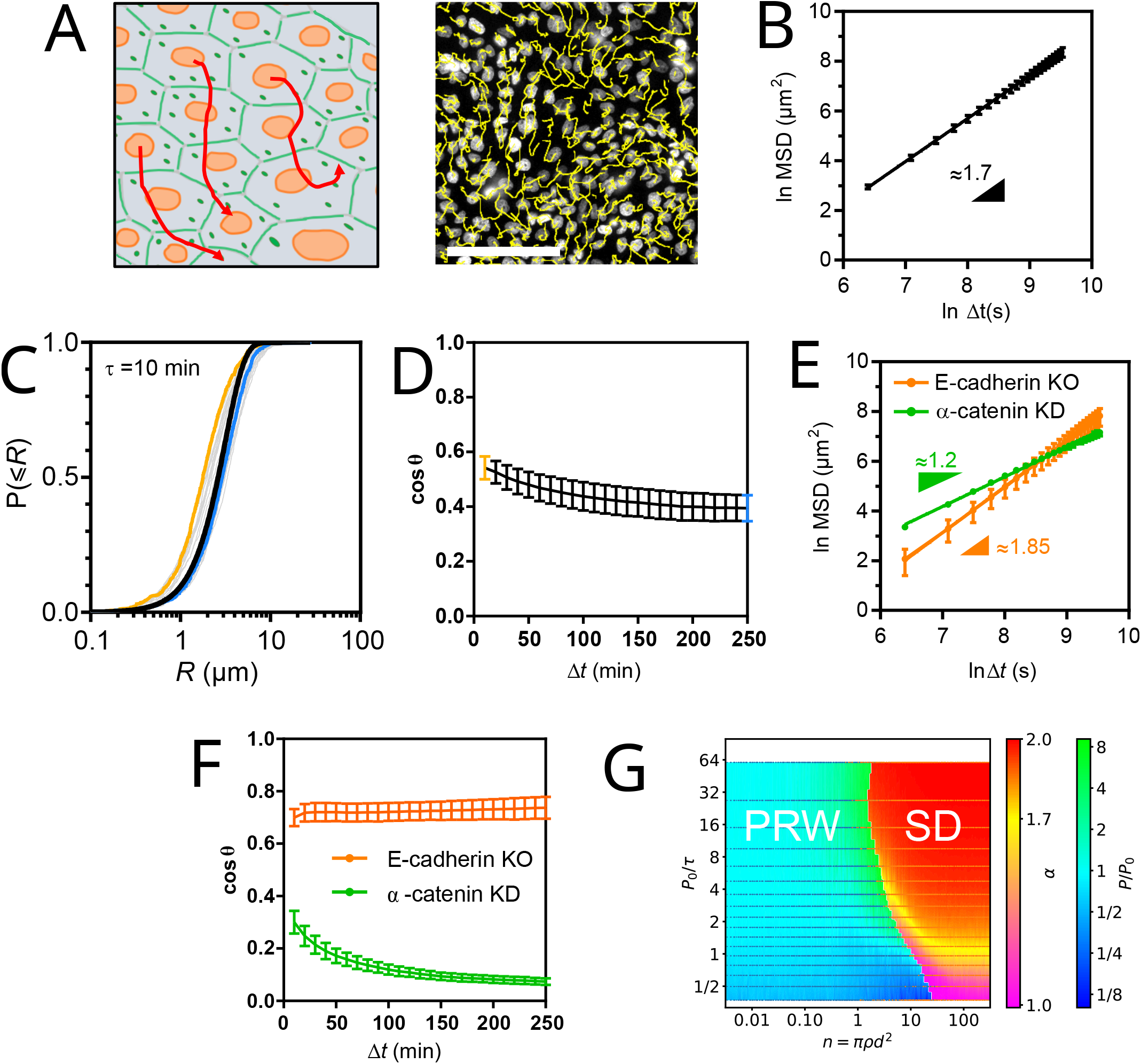
(A) Left: Schematic representation of the collective cell migration assay. Right: fluorescence image of nucleus-stained cells and their migration tracks in yellow. Scale bar = 100*µ*m. (B) MSD(Δ*t*) of WT cells fit with an anomalous diffusion model (straight line in ln-ln scale), the better model according to the Akaike information criterion. (C) Cumulative distributions of the step length *R* for *τ* = 10min. One gray curve per frame pair along a movie. The yellow and blue curves correspond to the first and last frames, respectively. The black line is a global fit of all blue curves with a cumulative Rayleigh distribution function. (D) Angular memory (cos *θ*(Δ*t*)) between Δ*t* = 10min (yellow dot) and Δ*t* = 250min (blue dot). The black line connects neighboring data points. (E) MSD(Δ*t*) of E-cadherin KO and *α*-catenin KD cells fit with a straight line in ln-ln scale. (F) Angular memory cos *θ*(Δ*t*) of E-cadherin KO and *α*-catenin KD cells. Data of (B-D) from 11 Fields Of View (FOV, 500-1000 tracks each) of 3 independent experiments. The colored lines connect neighboring data points. Data of (E-F) from 3 (E-cadherin KO) and 4 (*α*-catenin KD) FOV (500-100 tracks each) of 2 (E-cadherin KO) and 3 (*α*-catenin KD) independent experiments. Mean ± SEM. (G) Migration behavior of Self-Propelled Particles with Vicsek interactions as a function of *P*_0_ */τ* and number of interacting neighbors *n* = *πρd*^2^. Blue and orange points are simulation coordinates that resulted in persistent and super-diffusive behavior, respectively. Cold and warm color backgrounds are interpolations from points above of *P/P*_0_ and *α* values, respectively, and split the phase space in regions where persistent random walk (PRW) and super-diffusion (SD) occur, respectively (see Materials and Methods).

### The transition in directional memory emerges from Adherens Junction-mediated intercellular adhesion

To assess whether these effects resulted from simple steric hindrance due to crowding by neighboring cells or from genuine AJ-mediated intercellular adhesion, we examined the collective migration of cells depleted of *α*-catenin, without which cells cannot form AJs (54) (see Materials and Methods). Remarkably, the cells lost most of their super-diffusive behavior (with an anomalous exponent *α* dropping from 1.7 to ≈ 1.2) and recovered a higher cell speed, despite being as crowded as WT cells (Fig. 3E). Consistently, the angular memory vanished in the experimental time frame (Fig. 3F), as for single migrating cells. Next, to test whether cadherin subtypes could compensate for each other, we used cells depleted of E-cadherin but that retained cadherin-6 expression (54) (see Materials and Methods). These cells exhibited a slightly higher anomalous exponent than WT cells (Fig. 3E) and, consistently, long-term angular memory (Fig. 3F). This shows that E-cadherin specifically is largely dispensable, if not slightly detrimental, for cells to migrate in a super-diffusive manner, just as it is for the maintenance of AJs (54). Similarly, different cadherin subtypes can compensate for the defficiency of E-cadherin in other contexts (55– 57).

Therefore, the drop in cell speed and the transition in directional memory observed at confluence are the results of *bona fide* AJ-mediated intercellular adhesion, but independently of the cadherin subtype. Consequently, we define as a collective directional memory this long-term directional memory that arises from neighbor interactions.

### Velocity coordination between neighbors suffices to produce super-diffusion

Because velocity coordination is lost upon disruption of AJs by *α*-catenin depletion (Fig. S3E) (11), we sought to determine whether this effect of intercellular adhesion sufficed to confer a super-diffusive behavior to confluent cells regardless of other intercellular adhesion features. To do so, we turned to *in silico* simulations of interacting self-propelled particles (SPP, see Materials and Methods). First, we modeled individual cells as persistent random walkers with the same apparent speed and intrinsic persistence *P*_0_ in isolation for all cells, in compliance with single cell migration results. Accordingly, *P*_0_ is directly proportional to the time *τ* needed to make a step and a function of the intrinsic directional freedom of the cell between each step. Then, we modeled velocity coordination with a classic Vicsek interaction within a cell-scale distance *d* (58), and explored the MSD of particles at densities *ρ* covering isolated cells up to beyond confluence. We found that as the number of interacting neighbors *n* = *πρd*^2^ increased, the effective persistence *P* of the particles increasingly diverged from *P*_0_ before the migration behavior transitioned to superdiffusion (Fig. 3G). Strikingly, for a range of realistic values of interaction distance *d ≈* 10 *™* 30*µ*m, confluent density *ρ ≈* 2000 *™* 4000 cells*/*mm^2^ and various degrees of directional freedom affecting 1 ⪅ *P*_0_*/τ* ⪅ 10, the anomalous exponent fell around *α* ≈ 1.7 (yellow region), consistent with experimental observations (see Fig. 3).

Furthermore, this model predicted that provided a minimal number of interacting neighbors was reached, persistent random walkers would always transition to super-diffusers regardless of their intrinsic persistence. To test this, we tracked the single and collective cell migration of RPE-1 cells, another epithelial cell line (Fig. S3F, movies 4, 5). We found that single RPE-1 cell migration was also much better described by a persistent random walk than by a diffusion model, although with a persistence around 1 hr, making the difference hard to notice by eye within the experimental time window (Fig. S3E). Moreover, RPE-1 cell migration at confluence was again much better described by a super-diffusion model with an anomalous exponent around 1.6, as expected (Fig. S3E).

We conclude that velocity coordination is a feature of intercellular adhesion that suffices to produce super-diffusion at confluence in a manner that is very robust to the intrinsic persistence of cells, as predicted by our model.

### Molecular stabilization of Adherens Junctions by vinculin able to dimerize lowers cell migration speed

To further investigate the regulation of collective migration by intercellular adhesions, we examined the effects of confluence on vinculin mutant cell lines. All cell lines behaved as fluid tissues and their cells remained super-diffusive regardless of the expressed mutant (Fig. S4A, Fig. 4A, Table S3), showing that vinculin plays little or no role in the regulation of directional memory by intercellular adhesions.

**Fig 4.**
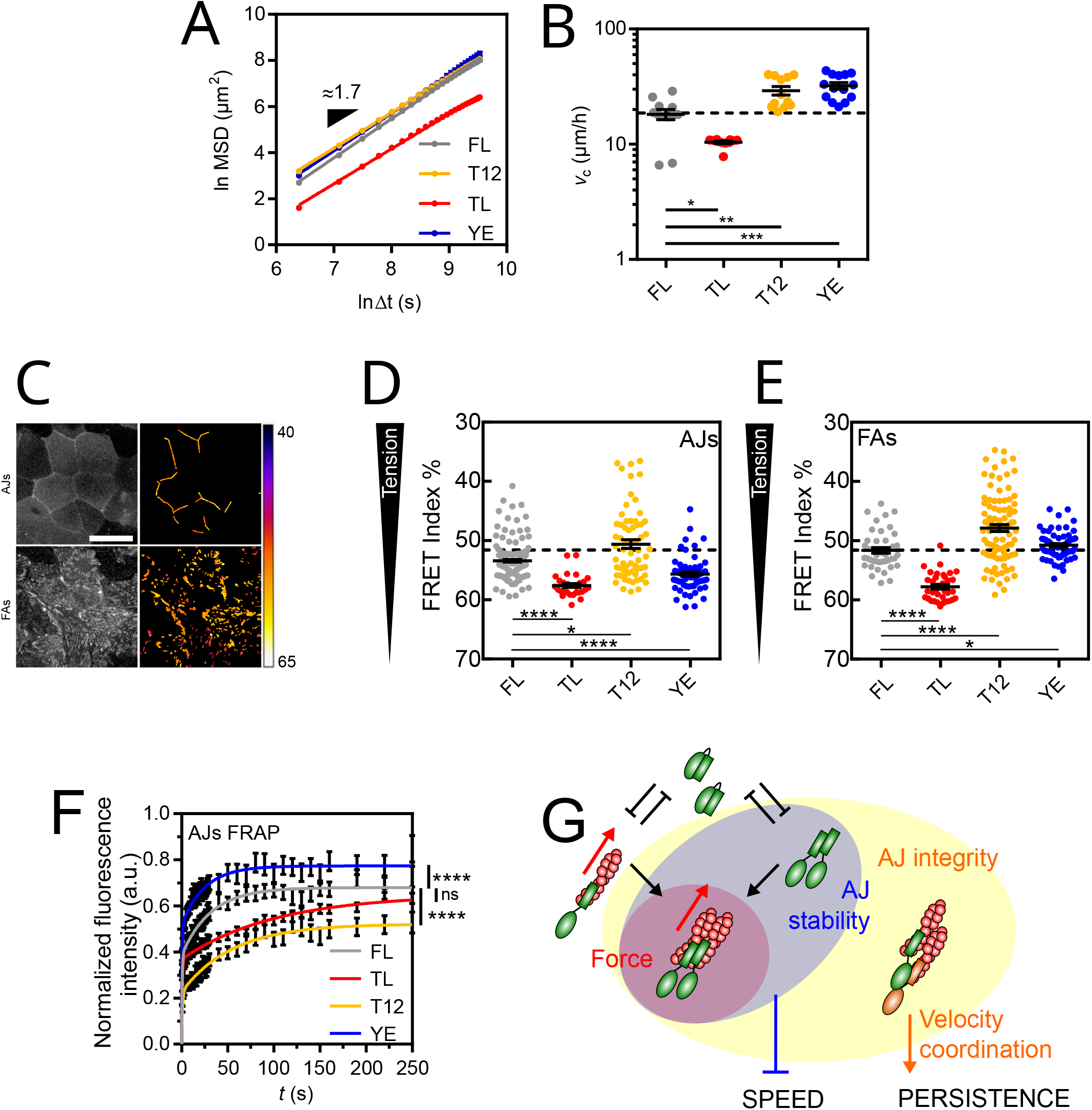
(A) MSD(Δ*t*) of vinculin FL and mutant cells fit with a straight line in ln-ln scale. (B) Apparent speed of vinculin FL and mutant cells at confluence. The dashed line indicates the mean of FL as a reference. (C) Left: direct fluorescence of vinculin TSMod in a confluent monolayer. Right: FRET Index map (see Materials and Methods). Top: in the AJs plane. Bottom: in the FAs plane. Scale bar = 20*µ*m. (D) FRET Index of TSMod in the FL, TL, T12 and YE constructs in AJs at confluence. The dashed line indicates the mean of FL as a reference. (E) FRET Index of TSMod in the FL, TL, T12 and YE constructs in FAs at confluence. The dashed line indicates the mean of FL as a reference. (F) Fluorescence recovery after photobleaching of FL, TL, T12 and YE constructs in AJs fit with a 2-phases association model. (G) Proposed Boolean model as a Venn diagram of the possible conformations and associated interactions of intact vinculin, and their individual and combined consequences on adhesion and migration: at AJs, *α*-catenin (orange) provides AJ integrity (connection to the cytoskeleton, yellow ellipse), which promotes persistence (from Fig. 3E,F). The head-tail interaction of vinculin (green) competes with both tail dimerization and actin filaments (red) binding. The ability to dimerize provides AJ molecular stability (blue ellipse, from panel F), which slows cell down (from Fig. S4C). AJ integrity and molecular stability are necessary but not sufficient for force transmission in AJs (red ellipse), the latter having no substantial influence on migration at confluence. Data of (A-B) from 12 (FL), 11 (TL, T12) and 14 (YE) FOV (500-1000 tracks each) of 3 independent experiments. Data of (D) from 169 (FL), 30 (TL) and 64 (T12, YE) contacts of 5 (FL) and 3 (TL, T12, YE) independent experiments, and (E) from 45 (FL), 35 (TL), 104 (T12) and 67 (YE) cells of 4 (FL) and 3 (TL, T12, YE) independent experiments. Data of (F) from 75 (FL), 137 (TL), 78 (T12) and 36 (YE) contacts of 4 (FL, TL, T12) and 3 (YE) independent experiments. Mean *±* SEM. Kruskal-Wallis and Dunn’s tests between each mutant and FL (B,D,E). Extra-sum-of-squares F test with shared immobile fraction between each mutant and FL as the null hypothesis (F). Benjamini-Krieger-Yekuteli-corrected p-values. *: p<0.05, **: p<0.01, ***: p<0.001, ****: p<0.0001.

Therefore, we examined cell speed. We reasoned that if vinculin neither plays a role in the regulation of cell speed by AJs, cell speed at confluence *v*_c_ can be entirely predicted from single cell speed *v*_s_ for each vinculin construct. Indeed, we found that the TL and T12 mutants remained slower and faster, respectively, than FL (Fig. 4B). Moreover, the three cell lines exhibited the same ≈ 10*µ*m*/*h decrease in speed as WT cells at confluence compared to single cells (Fig. S4C). This indicates that AJs decrease cell speed independently of the effects of vinculin mutations, which therefore have no other role in cell migration speed at confluence than those at FAs as in single cells. In contrast, the YE mutant was faster than FL at confluence (Fig. 4B), while it was slightly slower in single cells (see Fig. 2B). In fact, confluence induced a ≈ 10*µ*m*/*h increase, instead of a decrease in cell speed for this mutant (Fig. S4C). Altogether, this supports that functions impaired in YE, and not TL or T12, are key for the AJ-dependent decrease in cell speed at confluence.

At confluence, cortical actomyosin contractility promotes the recruitment and tension of vinculin in AJs (59). Here, all vinculin constructs colocalized with *α*-catenin and actin in intercellular contacts (Fig. S4D,E). To assess what distinguishes the YE mutant from the other constructs, we compared their relative recruitment and tension. In AJs, we found that YE showed a lower recruitment and tension than FL and T12, similar to TL, although to a milder extent (Fig. S4F, 4D). In FAs, in contrast, YE experienced a higher tension than FL, similar to T12, although to a milder extent (Fig. 4E). Altogether, the effect of YE on tension was neither different nor even stronger than that of the other mutants (Fig. S4G). Furthermore, there was no correlation between tension and cell migration speed among cell lines (Fig. S4H). This rules out tension as a regulator of collective cell migration speed.

Thus, we examined the stability of vinculin and its mutants by FRAP in AJs (Fig. S4I, movie 6), the molecular stability of which is based on that of vinculin (18). FRAP experiments showed that the YE mutant had a significantly smaller immobile fraction than FL vinculin, while the TL and T12 mutants had larger immobile fractions, just as in FAs, although only that of the T12 mutant was significantly different (Fig. 4F). Therefore, the lack of molecular stability in AJs is the unique difference between YE and the other mutants that appears at confluence.

Overall, these results support that the molecular stabilization of AJs (with TL, FL or T12), regardless of the tension they transmit through vinculin, equally lowers the speed of cell migration initially determined by FAs. In contrast, when AJs are unstable (YE), cells collectively migrate proportionally faster than single cells. Therefore, we propose that the ability of vinculin to dimerize ensures the molecular stability of AJs, which adapts the collective cell migration speed to the single cell migration speed (Fig. 4G). Consistently, we found that the optimal linear combination of changes in tension 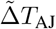 and stability 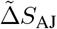 that could explain the change across vinculin constructs of the difference in cell speed between single cells and at confluence 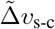 attributed all the weight to 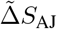. Furthermore, the dependence was better fit with a power law 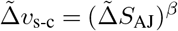 with *β* ≈ 0.2 (Fig. S4J, see Materials and Methods), reflecting how similar the effects of FL, TL and T12 on the difference in speed between single cells and at confluence were compared to the effect of YE.

### Super-diffusion overcomes the weak directional bias caused by adhesion-dependent sensitivity to external guidance cues

While cells adopt a super-diffusive behavior at confluence, they do so in the absence of any external guidance cue, and as a consequence without any preferred direction. Thus, we sought to evaluate how the addition of an external guidance cue would affect the spatial exploration efficiency of collectively migrating cells. In principle, a directional bias eventually overcomes any diffusion behavior on a sufficiently long time scale, resulting in a ballistic motion (MSD ∼ Δ*t*^2^, Δ*t* → ∞). To assess when it would occur, we wounded a confluent epithelial monolayer and tracked nucleus-stained cells as they invaded the free surface to close the wound (Fig. 5A, movie 7). Approximately 3 hours after wounding, the apparent speed of cells stabilized (Fig. S5A), consistent with previous reports (60), and was independent of the distance to the wound (Fig. S5B). Thus, we examined the MSD from 3hrs after wounding. Surprisingly, we found that the migration behavior remained super-diffusive with an anomalous exponent around 1.6 (Fig. 5B). Furthermore, as in the case of unbiased collective migration, the anomalous exponent decreased to around 1.2 for cells depleted of *α*-catenin, whereas cells depleted of E-cadherin remained superdiffusive, (Fig. 5C). Similarly, vinculin mutants did not affect super-diffusive behavior (Fig. 5D). Overall, these results show that space exploration efficiency remains dominated by the super-diffusive behavior of cells up to a few hours after exposure to an external guidance cue.

**Fig 5.**
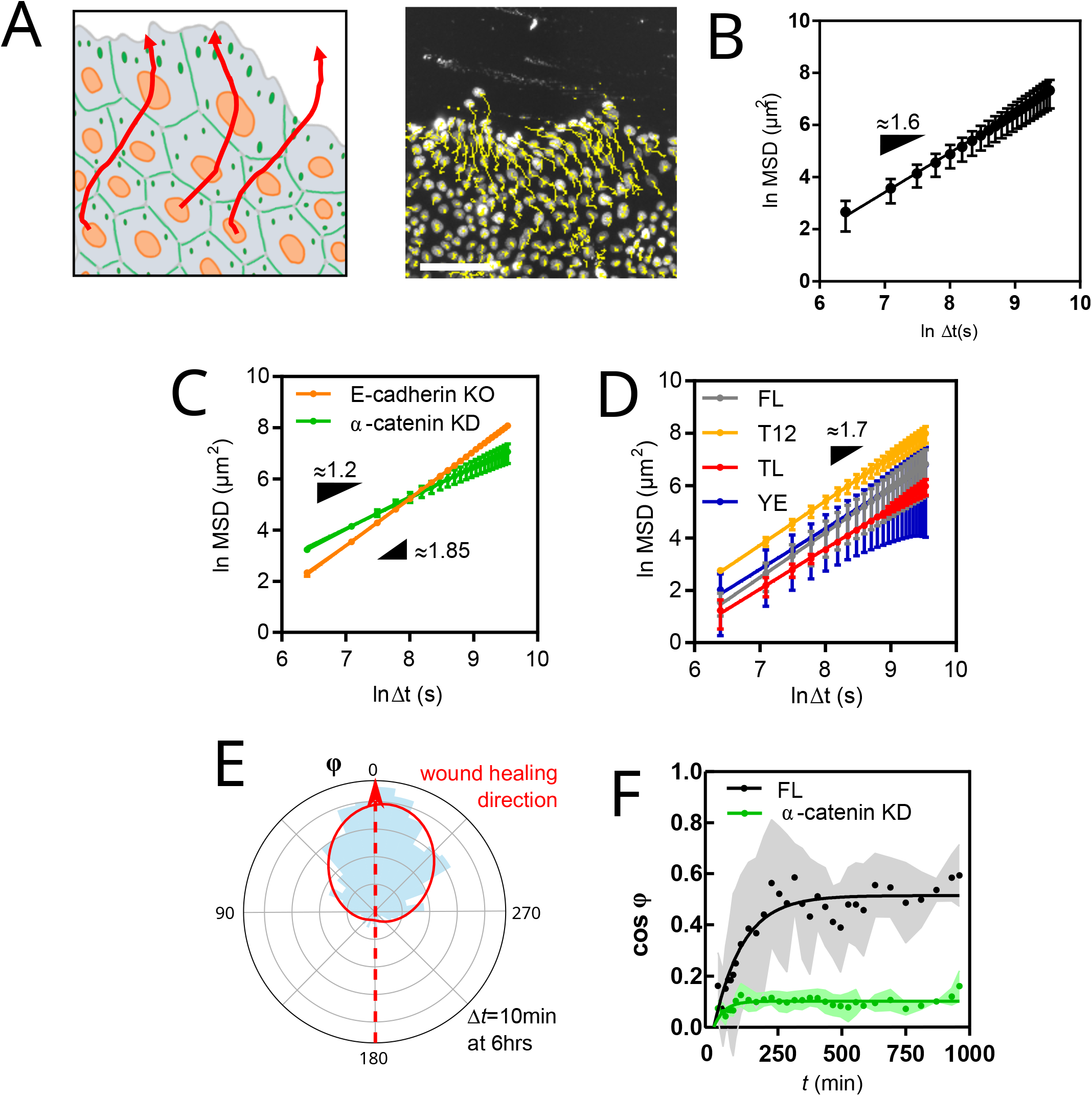
(A) Left: Schematic representation of the guided collective cell migration assay. Right: fluorescence image of nucleus-stained cells and their migration tracks in yellow. Scale bar = 100*µ*m. (B) MSD(Δ*t*) of WT cells fit with a straight line in ln-ln scale. (C) MSD(Δ*t*) of E-cadherin KO and *α*-catenin KD cells fit with a straight line in ln-ln scale. (D) MSD(Δ*t*) of vinculin FL and mutant cells fit with a straight line in ln-ln scale. (E) Angular distribution of apparent velocities of vinculin FL cells at 6hrs post-wound fitted with a Von Mises distribution giving a concentration *κ ≈* 1.3 (solid red line). (F) Directionality (cos *ϕ*(*t*), see Materials and Methods) of vinculin FL and *α*-catenin KD cells fit with one-phase association model. All data from 4 (all but YE) and 3 (YE) FOV (500-1000 tracks each) of 4 (all but YE and *α*-catenin KD) and 3 (YE, *α*-catenin KD) independent experiments. Mean *±* SEM.

This raised the question whether cell velocity was biased at all or if cells merely filled the wounded area by isotropic super-diffusion. To assess this, we examined the angular distribution of cell velocities. In the steady state speed regime (at 6hrs), a directional bias was clearly visible (Fig. 5E). The bias appeared within 3hrs post-wound and remained stable throughout (Fig. 5F). This shows that a directional bias of such magnitude maintained from 3hrs onward is too weak to substantially improve the spatial reach of collectively migrating cells in the time range of the experiment. Finally, we asked whether disrupted AJs would alter the sensitivity of cells to directional bias. Remarkably, cells depleted of *α*-catenin exhibited very little directional bias compared to WT cells (Fig. 5F). Together, this shows that AJ-mediated intercellular adhesion is required not only for cells to acquire long-term directional memory (see Fig. 3E, 5C), but also for them to perceive and migrate toward external guidance cues.

### Wound healing rate scaling with collective cell migration speed is linked to the ability of vinculin to dimerize by the regulation of Adherens Junction mechanosensitivity

Next, we wondered whether AJs were involved in the regulation of directed collective cell migration through additional mechanisms distinct from the regulation of directional memory and sensitivity to guidance cues by *α*-catenin (see Fig. 3E, 5C,F), and the regulation of speed by the ability of vinculin to dimerize and stabilize AJs components at confluence (see Fig. S4C). To do so, we assessed whether the rate of wound healing *v*_d_, at which the wounded epithelium covers the free surface (see Materials and Methods), could be predicted from the speed of collective cell migration at confluence in vinculin mutant cell lines (see Fig. 4B). We expected a correlation if no additional regulation depended on vinculin. We found that this was mostly the case, with the exception of the YE mutant. Indeed, while vinculin FL, T12 and TL cells closed the wound at a rate proportional to their migration speed at confluence, YE mutant cells were less efficient in wound healing than expected (Fig. 6A). This supports the idea that the ability of vinculin to dimerize, in contrast to other vinculin tail interactions, promotes wound healing in a manner that cannot be explained from its effects on collective migration at confluence, nor on single cell migration.

**Fig 6.**
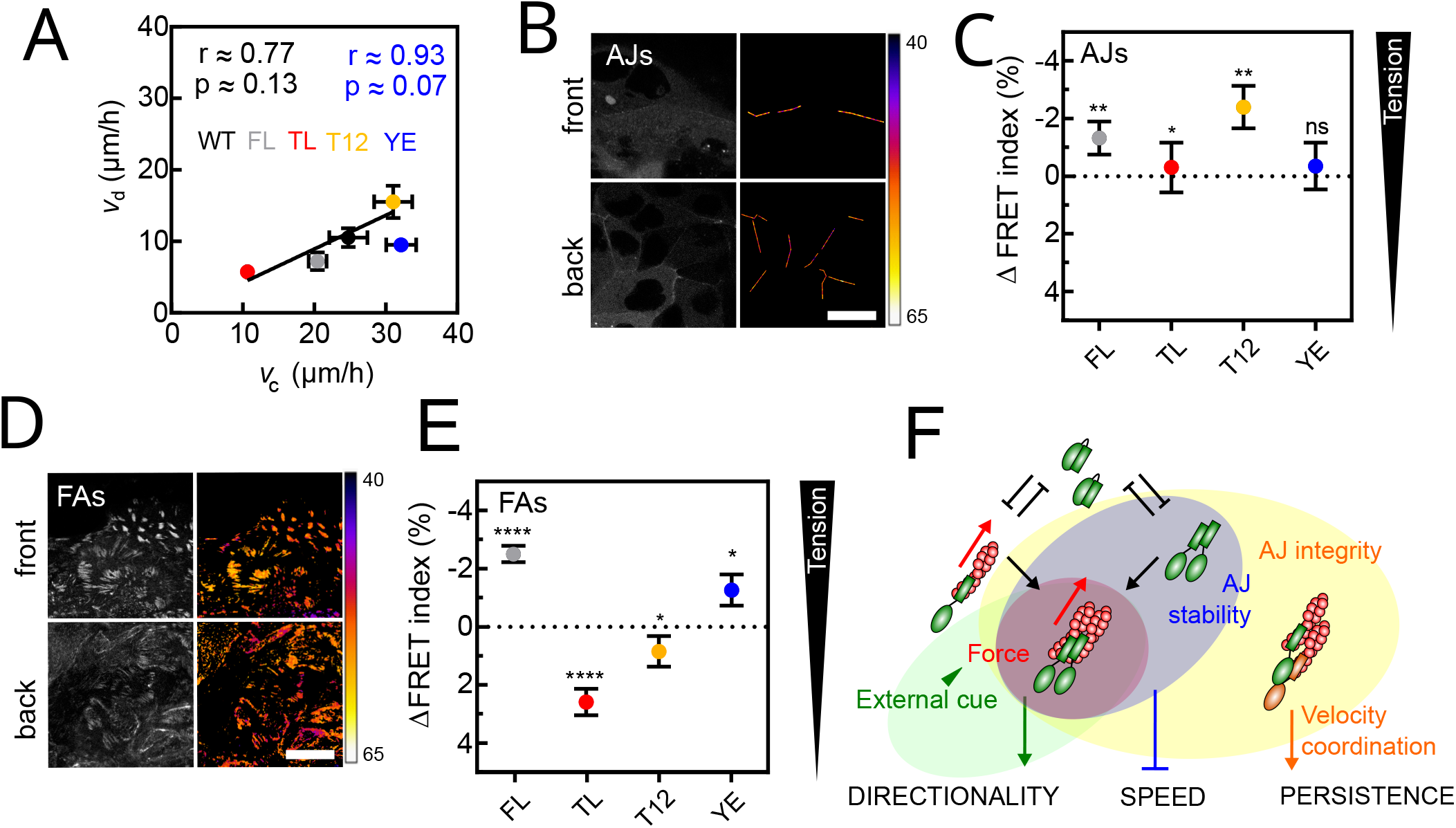
(A) Wound healing rate (see Materials and Methods) as a function of cell speed at confluence for WT, vinculin FL and mutant cells. Pearson coefficients r and p-values in black for all data, in blue minus YE. The black line is a linear fit without YE. (B) Left: direct fluorescence of vinculin TSMod in the AJs plane of cells at the wound front (top) and back (bottom). Right: corresponding FRET Index maps. (C) FRET Index difference between front and back cells at AJs of TSMod in vinculin FL and mutants. The dashed line indicates ΔFRET = 0 as a reference. (D) Left: direct fluorescence of vinculin TSMod in the FAs plane of cells at the wound front (top) and back (bottom). Right: corresponding FRET Index maps. (E) FRET Index difference between front and back cells at FAs of TSMod in vinculin FL and mutants. The dashed line indicates ΔFRET = 0 as a reference. (F) Proposed Boolean model as a Venn diagram of the possible conformations and associated interactions of intact vinculin, and their individual and combined consequences on adhesion and migration: at AJs, the ability of vinculin to dimerize and actin binding provide mechanosensitivity (red ellipse, from panel C) to stable AJs (blue ellipse, from Fig. 4F), which promotes directional migration in the presence of an external cue (such as the wound, green ellipse, from panel A). Data of (A) from 4 (all but YE) and 3 (YE) FOV of 4 (all but YE) and 3 (YE) independent experiments. Data of (C) from 55 (FL), 16 (TL), 56 (T12) and 26 (YE) front cells and 122 (FL), 30 (TL), 71 (T12) and 48 (YE) back cells of 4 (FL, T12) and 3 (TL, YE) independent experiments. Data of (E) from 144 (FL), 63 (TL), 124 (T12) and 46 (YE) front cells and 155 (FL), 84 (TL), 128 (T12) and 51 (YE) back cells of 7 (FL, T12), 4 (TL) and 3 (YE) independent experiments. Scale bars = 50*µ*m. Mean *±* SEM. Two-tailed Mann-Whitney test between front and back cells for each construct (C,E). Benjamini-Krieger-Yekuteli-corrected p-values. *: p<0.05, **: p<0.01, ***: p<0.001, ****: p<0.0001.

To determine how such an effect might emerge from this mutant, we therefore excluded its impact on adhesion molecular stability. Rather, we evaluated how vinculin tension perceived directed collective cell migration (Fig. 6B-E). Indeed, we had previously found that directed collective migration of epithelial cells imprints a transcellular gradient of molecular tension in E-cadherins, which are under higher tension away from the migration front (61). Consistently, we found that FL and T12 vinculin, both capable of dimerization and binding to actin, were increasingly recruited and under higher tension in AJs away from the migrating front (Fig. 6B, C). As expected, the TL mutant, incapable of dimerization or actin binding, showed minimal mechanosensitivity and, remarkably, the YE mutant only incapable of dimerization, did not show any (Fig. 6C). Therefore, actin binding is not sufficient for vinculin to exhibit a transcellular tension gradient in AJs in response to a directional cue, the ability to dimerize appears necessary.

In FAs (Fig. 6D, E), the TL mutant, which is insensitive to cytoskeletal tension by design, showed a lower FRET at the migration front. This suggests a cytoskeleton-independent increase in vinculin compression in back cells, as previously seen within single fibroblasts on adhesive patterns (62). The T12 mutant showed a shallow FRET gradient with slightly lower FRET at the front. Compared to TL, this suggests that an increase in cytoskeleton-dependent tension from front to back cells compensates for the opposite gradient of cytoskeleton-independent compression. FL vinculin showed a higher FRET at the migration front. This supports that the increase in tension from front to back cells overcompensates for the opposite gradient of cytoskeleton-independent compression. Remarkably, the YE mutant exhibited a mechanosensitivity similar to that of FL vinculin, with higher FRET at the front. This shows that dimerization is not required for vinculin to exhibit a transcellular gradient in FAs in response to a directional cue.

Together, this supports that the ability of vinculin to dimerize is involved in its mechanical sensitivity to external guidance cues only in AJs while actin binding is sufficient in FAs. Overall, this supports that the wound healing rate scales with collective migration speed as a function of the balance of mechanosensitivity to external guidance cues between AJs and FAs, a feature associated with the ability of vinculin to dimerize in AJs (Fig. 6F). Thus, we examined how the ratio between speed at confluence and wound healing rate 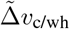 depended on the balance between mechanosensitivity in FAs 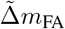 and in AJs 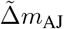, obtained from the FRET gradients between front and back cells in FAs and AJs, respectively (see Materials and Methods). Consistently, we found that 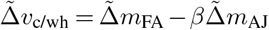 where *β ≈* 1*/*3 best described the changes across vinculin constructs (Fig. S5C), showing that a mechanosensitivity in AJs worth 1*/*3 that in FAs maintains the wound healing rate proportional to collective cell migration speed.

## Discussion

In this work, we studied the exploratory capacity of epithelial cells, the influence of cell interactions and guidance cues on this exploration, and the dynamics and mechanics of the adhesion complexes that cells use in this process. Remarkably, we show that intercellular adhesion with neighbor cells within the epithelium provides long-term directional memory for migration, as a result of velocity coordination. For this, proper interactions of AJs with the cytoskeleton through *α*-catenin are instrumental, but the type of cadherin involved is irrelevant. Additionally, cell speed depends on molecular stability and transmission of force in FAs and is impeded by the molecular stability of AJs, both of which are controlled by vinculin. Finally, our results support the idea of a specific role for the ability of vinculin to dimerize in AJs, which would regulate the mechanosensitivity of the epithelium as a whole to the guidance cue of a wound, and the wound healing rate of the epithelium.

Locomotion patterns of single cells have been observed and quantified for decades and can take many forms. When endowed with directional persistence arising from angular correlations between consecutive unit steps, cells adopt a so-called persistent random walk that eventually converges to a purely diffusive behavior, as initially observed in fibroblasts or single endothelial cells (4, 5). In contrast, in T cells and some metastatic cells that behave like Lévy walkers, the fat-tailed power-law distribution of step lengths results in super-diffusive behavior, even in the absence of temporal correlation in direction (7, 8). More recently, efforts have focused on developing analysis tools and models to account for heterogeneous migration properties along individual trajectories and among populations (63–67). Here, our results show that single epithelial cells undergo a persistent random walk, with persistence on the order of several minutes to an hour depending on the cell line, beyond which cells diffuse purely (Fig. 1). Thus, the persistence of epithelial cells is in the low range compared to endothelial cells, whose persistence time can reach several hours (5). Furthermore, comparing cell lines that express various forms of vinculin mutants, we observe that persistence does not correlate with cell speed, consistent with endothelial cells exposed to growth factors (5). Mechanistically, both cell speed and persistence are under the control of actin retrograde flow, via connections with cell-matrix adhesions for the former, and control of polarity factors for the latter (68). Our data are thus consistent with the idea that the abnormal connection of cell-matrix adhesions to the cytoskeleton via vinculin uncouples speed from persistence.

Previously, vinculin-KO fibroblasts or epithelial cells have been shown to migrate faster than their WT counterparts (22, 24, 29). Here, we show that the migration speed of epithelial cells decreases with the expression of FL vinculin even at low levels above endogeneous (Fig. 1). Overall, this is consistent with the idea that vinculin primarily acts as a migration moderator. Mechanistically, our results support that the head-tail interaction of vinculin challenges both force transmission and stability of vinculin in FAs, as has been observed in other cell types (30, 44, 47–50), and is countered by the actin-vinculin tail interaction, which provides force transmission but is unnecessary for stability, and the ability of the tail to dimerize, which associates with higher stability and is dispensable for force transmission (Fig. 2D,E). As a result of this feedback, stability and force transmission can be further promoted by the interaction of the vinculin tail with actin and its ability to dimerize, respectively, indirectly through their antagonism with the head-tail interaction of vinculin. Besides, a previously evidenced catch bond between the tail of vinculin and actin provides an additional mechanism for stabilization of vinculin in adhesions (69). Altogether, the combination of maximum force transmission and stability predicts maximum cell speed (Fig. 2B). Therefore, we propose that the speed of single cells results from the combination of force transmission and stability in FAs (Fig. 2F, S2I).

At confluence, we show that epithelial cells adopt a super-diffusive behavior that a persistent random walk cannot explain (Fig. 2). This behavior has previously been observed in single cells or in a collective for a variety of cell types (6–8, 13–15). However, proposed models involved non-Gaussian displacement distributions, which are sufficient to support super-diffusion without long-term memory. Moreover, whether super-diffusion in the collective emerged from intercellular interactions or from cells that already walk alone in this manner remained to be demonstrated. Here, we show that super-diffusion can be explained by a pure fractional Brownian motion with Gaussian distribution of displacement and long-term directional memory, which emerges from intercellular interactions (Fig. 2A-D).

Previously, sub-diffusive fractional Brownian motion has explained subcellular particle mobility in the crowded cytoplasm through interactions with the viscoelastic medium (70). In contrast, we show here that a strong super-diffusive fractional Brownian motion emerges from intercellular adhesion mediated by AJs bound to the cytoskeleton but independent of cadherin type, and not just steric hindrance due to the mere presence of neighboring cells (Fig. 3E,F). Remarkably, the same AJ-based mechanism also allows for velocity coordination between neighboring cells (Fig. S3E) (11). The interplay between persistent migration and elastic coupling provided by AJs was previously shown to be sufficient for velocity coordination (71). We further show with our minimal agent-based model that such coordination is sufficient to explain the transition to super-diffusion (Fig. 3G). Remarkably, the transition occurs in a region where *α* ≈ 1.7 for a number of interacting neighbors *n* ≈ 6, about the number of direct neighbors at confluence, and *P*_0_*/τ* ≈ 3. This allows us to determine the decision time *τ* , which is intrinsic to persistent random walks, but cannot be experimentally determined from single cell migration experiments that only reveal *P*_0_. For MDCK cells with persistence *P*_0_ ≈ 10min, this corresponds to a decision time *τ* of a couple minutes. For RPE-1 cells with persistence *P*_0_ ≈ 1hr, *τ* would rather be several tens of minutes to an hour. This suggests that RPE-1 cells are more persistent due to their longer decision time, rather than narrower directional freedom. Discovering the respective molecular determinants of decision time and directional freedom appears as an exciting future research direction. In the same modeling context, others have shown that velocity coordination also results in spontaneous fluctuations in particle density (72). We thus propose that at confluence, such spatiotemporal fluctuations in the number of interacting neighbors generate variations in instantaneous persistence of cells along their trajectories, ultimately resulting in a distribution of persistence times across time scales. Consistently, the MSD of a persistent random walk at time *P* scales as *P* ^*e−*1^, the anomalous exponent being about that of the super-diffusive MSD, as if super-diffusing cells experienced a distribution of persistence times *P* that spanned all our experimentally accessible time scales.

In addition to inducing a transition from short-to long-term directional memory, we show that AJs have a second role: they globally decrease the apparent speed of cells, provided that vinculin is stable at AJs. Indeed, perturbation of the ability of vinculin to dimerize leads to its low stability in AJs and faster than expected cell migration (Fig. 4B,F, S4C). Remarkably, this increase in cell speed is not the result of relaxed constraints on cortical deformations, as seen previously in unjamming transitions (73, 74), since vinculin mutants all exhibit similar shapes indicative of a fluid phase (Fig. S4A). Alternatively, we speculate that the destabilization of AJ components results in more complex changes, possibly due to the impaired ability to bundle actin of the vinculin mutant (43). Interestingly, a recent study has evidenced how mechanosensitive AJs downregulate lamellipodial activity in collectively migrating cells (75). However, we have previously shown using a minimal reconstitution approach that the down regulation of lamellipodial activity by AJs does not decrease the migration speed of cells (11), but desmosomes do (76). This suggests that the ability of vinculin to bundle actin might indirectly regulate desmosome formation by AJs. Finally, the possibility remains that the stability of vinculin in AJs indirectly alter FAs, and thereby migration speed, in ways that we have not examined. Future studies may address these hypotheses.

Overall, intercellular adhesion normally impairs apparent cell speed but induces long-term directional persistence. As a result, while cells are collectively less efficient than single cells in exploring space at time scales of less than 3hrs, they can become more efficient beyond that (Fig. S3B). Thus, this emergence of long-term collective persistence may have a profound impact on tissue morphogenesis, regeneration, and homeostasis in a variety of contexts. In the case of wound healing, cells are further exposed to an external guidance cue. Here we show that this signal is sufficient to bias the velocity of each cell at the smallest experimental time scale (Fig. 5E). Detection of this signal is an emergent property of the collective, as disrupting intercellular adhesion renders cells nearly blind to the free space to invade (Fig. 5F). This is reminiscent of collective durotaxis and chemotaxis (77, 78). However, we show that this directional bias remains too small to drive ballistic motion in less than the longer experimental time range, such that cells remain super-diffusive throughout (Fig. 5A-D). We expect the directional bias to lead to more ballistic cell motion at longer time scales, with a lower limit of at least 3hrs.

This third role of AJs in directing collective cell migration is also dependent on vinculin. Specifically, wound healing rates scale with collective cell migration speeds when the ability of vinculin to dimerize is preserved (Fig. 6A). In this context, we propose that the ability of vinculin to dimerize is required to provide mechanosensitivity to AJs, so that both AJs and FAs can transmit molecular tension gradients across the tissue (Fig. 6B-E). This is again reminiscent of the requirements for supracellular durotaxis, which arises from intercellular force transmission (77). We speculate that the ability of cells to stiffen under force in a way that depends on the ability of vinculin to dimerize may be involved (43).

Whether vinculin dimerization, or another unknown independent function disabled by the YE mutation or phosphorylation of Y1065, is specifically involved in the adhesion and migration phenotypes described above may be addressed in future studies. Likewise, whether the effects are stronger in the absence of endogenous vinculin may be addressed in a null background. Nevertheless, intermolecular FRET is compatible with the existence of vinculin dimers in FAs (40), and the ability of such dimer to antagonize vinculin head-tail interaction and to promote actin bundling remains the most parsimonious mechanism that can explain the stabilization of vinculin in FAs and the stiffening of cells described here and elsewhere (43, 79). Interestingly, a recent study showed that another nearby phospho-mutation that relaxes vinculin tension and favors its closed-conformation in FAs and AJs also alters directed collective cell migration (80). Whether this mutant affects cell migration through FAs, AJs, or both remains to be determined.

In summary, we conclude that the spatial exploration efficiency of guided collective migration depends on the scale considered: at short time scales (<3hrs for MDCK cells), the intrinsic migration speed of individual cells is the limit, which can be overcome by collective directional memory at intermediate time scales (>3hrs for MDCK cells), a tissue-level property that guidance cues may additionally enhance, but at much longer time scales (»3hrs) (Fig. S5E). We speculate that this generic framework applies qualitatively to guided collective migration in general and hope that future studies will test this prediction.

## Materials and Methods

Live MDCK type II G cells stably expressing fluorescently tagged proteins were monitored on a wide-field epifluorescence inverted microscope for migration experiments, on a scanning spectral confocal microscope for FRET experiments, or on a spinning disk confocal microscope for FRAP experiments. Image analyzes were performed with Imaris and ImageJ, simulations with R, data analyzes with Matlab and R, and statistics with Prism, R and Python. Complete materials and methods are available in Supplementary Materials and Methods. New materials, reagents, and simulation and analysis scripts are available upon request.

## Data availability

This study does not include data deposited in external repositories.

## ACKNOWLEDGEMENTS

We thank the members of the laboratory, C. Leclainche, S. Etienne-Manneville, B. Ladoux, P. Ronceray and R. Voituriez for insightful discussions. We thank C. Grashoff (Westfälische Wilhelms-Universität Münster), B. Ladoux and R.-M. Mège (Institut Jacques Monod), W. James Nelson (Stanford U.) and K. Schauer (Institut Gustave Roussy) for the gift of plasmids and cell lines. This work was supported in part by the Centre national de la recherche scientifique (CNRS), the French National Research Agency (ANR) grants (ANR-17-CE13-0013, ANR-17-CE09-0019, ANR-18-CE13-0008, ANR-21-CE13-0048), the Investments for the Future program of the French Government (LabEx Who am I? ANR-11-LABX-0071, Université Paris Cité ANR-18-IDEX-0001) and the Association pour la Recherche contre le Cancer (ARC-PJA22020060002255). We acknowledge the ImagoSeine facility, member of the France BioImaging infrastructure (ANR-10-INSB-04), and the Institut Jacques Monod IT team. HC was supported by La Ligue contre le Cancer (TAYS18872) and Fondation Recherche Médicale (FDT202001010843).

