## Supplementary material for "Collective directional memory controls the range of epithelial cell migration": SI materials methods and figures (from another version of this preprint: version 4, 2026-08-04)

### Materials and Methods

**Cells.** MDCK type II G dog cells were maintained in low glucose DMEM with phenol red supplemented with 10% v/v Fetal Bovine Serum (FBS) and Penicillin 10 U/mL + Streptomycin 10 µg/mL, and hTERT RPE-1 human cells in DMEM/F12 with 10% v/v FBS, at 37°C and 5% CO2. Cells were regularly tested for mycoplasma contamination. Cells were transfected with Turbofect transfection reagent (Thermo Scientific) according to the manufacturer's instructions in a WT background to avoid intermolecular FRET between exogenous constructs. 24 hours post-transfection, cells were cultured under drug selection (G418 sulfate 200 µg/mL) for 15 days and FACSed for intermediate expression to avoid level-dependent ectopic localization of exogenous constructs. The generated stable lines were cultured under drug selection. MDCK WT and $\alpha$-catenin KD cell lines were kind gifts from the laboratory of W. James Nelson (Stanford U.) (1). The E-cadherin KO cell line was a kind gift from the laboratory of B. Ladoux and R. M. Mège (Institut Jacques Monod) (2). The RPE-1 cell line was a king gift from the laboratory of K. Schauer (Institut Gustave Roussy). Cell lines were not authenticated. Cells were imaged at 37°C and 5% CO2 in Fluorobrite medium supplemented with 10% v/v FBS, Penicillin 10 U/mL + Streptomycin 10 µg/mL, 2.5 mM L-Glutamine and 20mM HEPES on 1.0 borosilicate, 18 or 32mm Ø round glass coverslips or on 2- or 4-well Nunc™ Lab-Tek™ Chambered Coverglass supports (Thermo Scientific) previously coated with collagen IV by incubation with a solution (50 µg/mL in 0.5M acetic acid) of collagen from human placenta Bornstein and Traub Type IV (Sigma) for 10 minutes before air-drying.

**Plasmids.** VinculinTS (addgene #26019) and Tailless VinculinTS (#26020) were a kind gift of the laboratory of Carsten Grashoff (Westfälische Wilhelms-Universität Münster) (3). The following mutations were introduced with the QuikChange II XL Site-Directed Mutagenesis Kit (Agilent Technologies):
T12 (5'-GAGGTGGCCAAGCAATGTACTGTCGCAGCCA
TTGCAACAAACCTC-3', 5'-CTCACAGACCTGTAAGA
GGTTTGTTGCAATGGCTGCAGCAGTACA-3'),
Y1065E (5'-ATCTGCAGAATTCTTACTGCTCCCATGGG
GTCTTTCTGACC-3', 5'-GGTCAGAAAGACCCCATGG
GAGCAGTAAGAATTCTGCAGAT-3'),

**Migration assays.** For single cell migration, 36 hours before imaging, cells were passaged at a 1:10 dilution with vigorous pipetting, then 4 to 6 hours before imaging cells were detached once again, resuspended with vigorous pipetting and plated at about 6 cells/mm². For collective migration, 24 hrs before imaging, cells were resuspended and plated at about 600 cells/mm². For biased collective migration, cells were additionally incubated at 37°C in DPBS for 5 to 10 minutes before imaging to weaken calcium-dependent cell-cell contacts and a straight-lined scratch was made in the monolayer using a 200 µL pipette tip. DPBS was then replaced with imaging medium. Prior to imaging, nuclei were stained by incubation in Hoechst 0.1 µg/µl (Thermo Scientific) imaging medium for 20 minutes followed by further dilution to 0.05 µg/µl. Imaging was performed on a ZEISS Axio Observer Z1 widefield microscope with 20x Plan-Apochromat/0.8 dry objective. Hoechst fluorescence was imaged by excitation with a CoolLED pE-300 UV fluorescent lamp, excitation and emission filters 335-383 and 420-470, and an ORCA-Flash4.0 LT camera. Multi-position and multi-tile images were acquired every 10 minutes overnight.

**Migration analyses.** Nuclear staining detection, cell segmentation and track generation were performed with Imaris 9.2 for single cell migration and Fiji with custom scripts based on Stardist and TrackMate plugins (4, 5) for collective and biased collective migration. Visual inspection of tracks against raw movies enabled manual correction if necessary. Further data analyses were performed with Matlab with custom scripts and @msdanalyser (6). The apparent cell speed $v$ is defined as the distance traveled during the smallest time interval (10 min) and is obtained directly from the cell tracking in Imaris and TrackMate. The speed is averaged over each track, then over the cell population for single cell migration experiments. The speed is averaged over each field of view for each time point and over the duration of the experiment for collective and biased collective migration. The mean square displacement (MSD) is: $\text{MSD}(\Delta t) = \langle(\Delta \mathbf{r}_i(\Delta t))^2\rangle = \langle(\mathbf{r}_i(t+\Delta t) - \mathbf{r}_i(t))^2\rangle$, where $\mathbf{r}_i$ is the position of the cell $i$ at time $t$, $\Delta t$ the lag time, and $\langle\rangle$ is the average for all cells and $t$. For each experiment comprising multiple field of view acquisitions, a weighed mean MSD was calculated to correct for differences in track duration. For each cell line and condition, the average MSD was calculated as a weighed mean between experimental replicates to account for potential differences in experiment duration. $\text{MSD}(\Delta t)$ curves were fit with the Fürth equation $\text{MSD} = 4D(\Delta t - P(1 - \exp(-\Delta t/P)))$ where $D$ is the coefficient of diffusion and $P$ the persistence time, as a model of persistent random walk, or with $\text{MSD} \sim \Delta t^{\alpha}$ (straight line of slope $\alpha$ in $\ln - \ln$ scale), as a model of anomalous diffusion, with anomalous exponent $\alpha > 1$ for super-diffusion. The step length $R_{i,t}(\tau)$ of a cell $i$ is $\sqrt{(\mathbf{r}_i(t+\tau) - \mathbf{r}_i(t))^2}$, where $\tau$ is the step duration. Cumulative distributions of step lengths are displayed for all cells at every time point. Curves from all cells and all time points were fit together with the cumulative distribution function of the Rayleigh distribution $\text{P}(< R) = 1 - \exp -R^2/2\sigma^2$ where $\sigma$ is the scale parameter. The angular memory $M_{i,t}(\Delta t)$ is defined as the cosine of the angle $\theta_{i,t}$ between consecutive step vectors $\mathbf{L}^{t}_{i,t-\Delta t}$ between times $t - \Delta t$ and $t$ and $\mathbf{L}^{t+10\text{min}}_{i,t}$ between times $t$ and $t + 10\text{min}$ of a cell $i$ at time $t$, such that the step duration $\tau$ of the first vector and lag time $\Delta t$ before the next are equal. $M_{i,t}(\Delta t) = \cos\theta_{i,t}(\Delta t) = \cos(\mathbf{L}^{t+10\text{min}}_{i,t}, \mathbf{L}^{t}_{i,t-\Delta t})$. Figures display $\langle M_{i,t}(\Delta t)\rangle$. This metric measures the duration of the past that the cell considers when deciding on its direction. It decays to 0 when $\Delta t \to \infty$ for persistent random walkers and remains constant for fractional Brownian motion, similarly to the auto-covariance function of fractional Brownian motion $C(\Delta t/\tau) = ((\Delta t/\tau+1)^{\alpha} + |\Delta t/\tau - 1|^{\alpha} - 2(\Delta t/\tau)^{\alpha})/2$, which is a constant $C = 2^{\alpha-1} - 1$ for all $\Delta t = \tau$, as in our definition of angular memory (while it decays to 0 when $\Delta t/\tau \to \infty$) (7). Ballistic motion yields 1 whatever $\Delta t$ and uncorrelated random walks, whether anomalous or not, yield 0. For this reason, and because $M_{i,t}(\Delta t)$ was found more robust to noise than $C(\Delta t/\tau)$, we considered this metric a more compelling discriminant between fractional Brownian motion and other uncorrelated super-diffusive walks. The cell shape index is defined as $q = p/\sqrt{A}$, where $p$ and $A$ are the perimeters and areas of the cells approximated by a Voronoi tessellation around nucleus centers, as done previously (8). Coordination is defined as the cosine of the angle $\gamma$ between the apparent velocities of cell pairs distant of $d$. Directionality is defined as the cosine of the angle $\phi$ between the cell apparent velocity vector and the direction orthogonal to the wound. Figures display the population average and standard error. Curves were fit with a one-phase association equation: $\cos\phi = \cos\phi_{\text{max}}(1-\exp(-t/\tau_{\text{d}}))$ where $\cos\phi_{\text{max}}$ is the average directionality at the infinite time limit and $\tau_{\text{d}}$ the characteristic time to reach it. Wound closure speed is defined as the rate of wound half-area $A/2$ decrease in $\Delta t = 30$min normalized to wound length $l$: $\text{WC} = (A(t+\Delta t) - A(t))/(2l\Delta t)$.

**Simulations.** Self Propelled Particles (SPP) simulations were performed in R software (version 4.2.2) on $N = 200$ particles within a 2D square box of side $L$ with periodic boundary conditions for $T = 600$ steps of duration $\tau$. From an initial condition of random positions and orientations of all particles, the position of each particle $i$ is updated according to $\mathbf{r}_i(t+\tau) = \mathbf{r}_i(t) + \mathbf{v}_i(t+\tau)\tau$, where $\mathbf{v}_i$ is the velocity of particle $i$ with constant modulus $v$ and an orientation $\theta_i(t+\tau) = \arg\left[\sum_{j,|\mathbf{r}_i(t)-\mathbf{r}_j(t)|<d} \mathbf{v}_j(t)\right] + \eta\xi_i$. The first term accounts for an interaction that causes velocity alignment between neighbors within a distance $d$ (9). The second term is a white noise $\xi_i(t)$ over $[-\pi,\pi]$ bounded by $\eta \in [0,1]$, which defines the directional freedom of an isolated particle. The intrinsic cell persistence, i.e. that of an isolated cell, directly relates to $\eta$ and $\tau$ through $P_0 = \tau(\overline{\cos\theta_i})^{1/2}/(1-\overline{\cos\theta_i})$ where $\overline{\cos\theta_i} = \sin\eta\pi/(\eta\pi)$, by generalization of Fürth's approach (10) to two dimensions. The apparent speed $v = 2.5$µm.min$^{-1}$ was fixed in the typical range of experimental measurements. The total particle density $\rho = N/L$ was set to various values within $[10^{-5}, 10^{-2}]$cell.µm$^{-2}$ to match single ($\rho \ll 10^{-3}$cell.µm$^{-2}$) or collective ($\rho \geq 10^{-3}$cell.µm$^{-2}$) cell migration assays. The interaction distance $d = 5-40$µm was varied around the cell size scale. The directional freedom $\eta$ was varied within $]0,1[$ for the sake of exhaustiveness. Simulated tracks were analysed as live cell tracks to compute the MSD($\Delta t$) that was fitted in ln-ln scale with the minpack.lm R package. Fit parameters $P$ and $\alpha$ were averaged over 5 simulations for each combination of parameter values and linearly interpolated on a 400x400 grid between extreme values of $P_0/\tau$ and $\pi\rho d^2$ for representation purposes. The $P-\alpha$ boundary was set at the transition in majority behavior within moving boxes of size 0.25 in log-log scale.

**Live confocal and molecular tension microscopy.** Cells were imaged on a ZEISS LSM780 confocal microscope with a 63x/1.4NA oil immersion objective. The mTFP1 donor fluorophore was excited by a 458 nm Argon laser line. Fluorescence emission was detected in the 473nm-561nm spectrum with a spectral resolution of 8.8nm (10 channels) by a GaAsP (Gallium Arsenide Phosphide) detector. The pixel size was ≈0.09 x 0.09 x 0.37µm. Z-stacks of 10-15 steps were acquired. Image analysis was performed in Fiji. Segmentation of FAs on the first slice of the z-stack was performed by a custom macro that subtracts the background, applies a Gaussian filter, thresholds, localizes particles of an area of 0.7 µm² or more and a circularity factor between 0 and 0.95 and displays the final Region Of Interest (ROI) set with the HiLo look up table to enable manual removal of any spurious or saturated ROI. Segmentation of AJs was performed manually on other slices of the z-stack. Intensity in AJs is normalized against mean intensity in FAs of the same cell after subtraction of the background measured in a nearby cytoplasmic region for every slice of the z-stack. The FRET index $I_{\text{F}} = I_{\text{FRET}}/(I_{\text{donor}} + I_{\text{FRET}})$ is calculated using the PixFRET plugin in Fiji (11) from the intensities $I$ of the channels around 494 (donor) and 530nm (FRET) of the spectral stack, with a 2px radius Gaussian filter and a threshold above 1x mean background intensity. Using the ROI set generated from the corresponding intensity image, the ROI pixels belonging to one cell are combined to obtain an average FRET index value of FAs or AJs per cell. By design of the TSMod sensor inserted into the vinculin constructs, a decrease in FRET index reports an increase in tension per vinculin (3), averaged over all proteins within the ROI.

**Fluorescence recovery after photobleaching.** FRAP was performed on a Yokogawa Spinning Disk CSU-X1 microscope. Cells were imaged with a 63x/1.4NA oil immersion objective. Fluorescence was acquired by 491nm laser excitation with a GFP 525/50 emission filter and a sCMOS PRIME 95 camera (Teledyne-Photometrics). Photobleaching was achieved by a 476nm FRAP laser. Images were analyzed with ImageJ. Fluorescence in ROIs was background-subtracted, corrected for photobleaching due to imaging and normalized by the fluorescence before photobleaching. The lack of contrast between photobleached adhesion structures and their surrounding at $t = 0$ ensured 100% photobleaching. The average curves for each condition were then fitted with a two-phase association equation: $F = F_{\text{fast}}(1-\exp(-t/\tau_{\text{fast}})) + F_{\text{slow}}(1-\exp(-t/\tau_{\text{slow}}))$ where $F_{\text{fast}}$ is the fluorescence of the mobile fraction characterized by the time $\tau_{\text{fast}}$ that accounts for the very fast initial recovery and $F_{\text{slow}}$ the fluorescence of the mobile fraction characterized by the time $\tau_{\text{slow}}$ that accounts for the second recovery phase. The immobile fraction is therefore $F_{\text{im}} = 1 - F_{\text{fast}} - F_{\text{slow}}$. We use the immobile fraction of vinculin and its mutants as a measure of vinculin stability and, by extension of the molecular stability of adhesions, because the stability of other adhesion proteins is regulated by the stability of vinculin (12–14).

**Immunofluorescence.** Cells were seeded on 18 mm diameter coverslips. Cells were washed with 0.5 mM Ca2Cl in PBS, fixed with 4% paraformaldehyde (PFA) in the washing medium for 5 min at room temperature, washed again with PBS 3 times for 5 min each, permeabilized with 0.5% triton in PBS for 5 min at room temperature, washed with PBS, incubated for 5 min at room temperature with 50 mM NH4Cl in PBS, incubated for 30 min in Blocking Buffer (BB, 1% Bovine Serum Albumin, 1% donkey and goat sera in PBS), incubated with mouse 15D9 anti $\alpha$-catenin primary antibody (1:200, sc-65479 Santa Cruz Biotechnology) and AlexaFluor568-phalloidin (1:200, Invitrogen) in BB for 1 hour in the dark, washed with PBS 3 times for 5 min each, incubated with anti mouse DyLight 633 secondary antibody (1:200, Invitrogen) in BB for 45 minutes in the dark, washed with PBS 3 times for 5 min each, mounted on Superfrost slides (Thermo Scientific) in Fluoromount (Sigma) and let dry in the dark overnight. Imaging was performed on a ZEISS LSM780 confocal microscope with a 63x/1.4NA oil immersion objective. The vinculin constructs, AlexaFluor 568 and DyLight633 were excited through a 488/661/633 mirror beam splitter by a 488 nm Argon laser line, a 561 nm Diode laser, and a HeNe 633 nm laser, respectively. Fluorescence emission was detected in the 499-534, 571-618 and 651-696 ranges, respectively, by a GaAsP and 2 standard alkali PMT (photo multiplier tube) detectors. The pixel size was ≈0.09 x 0.09 x 0.37µm. Z-stacks of 10-15 steps were acquired. Image analysis was performed in Fiji. Intensities were measured along 4 pixels-wide lines on maximum intensity projections of z-stacks excluding slices where FAs were visible.

**Western Blot.** Cells were washed with PBS at 4°C, lysed on ice in RIPA buffer (with protease and phosphatase inhibitors) by scraping, collected in centrifugation tubes, vortexed for 1 minute and centrifuged at 13000 rpm for 15 minutes at 4°C. Supernatant was then collected and diluted with an equal volume of Laemmli Buffer 2x (Sigma). Samples were loaded on a Mini-Protean TGX Precast gel (BioRad), ran for 1 hour at 100 V in running buffer (Tris-Glycine buffer 1x, SDS 0.005%) and transferred onto a nitrocellulose membrane in a 10% methanol transfer buffer (Tris-Glycine 1x, methanol 10%) for 1 hour at 100 V. The transfer was verified by staining with Ponceau S solution. The membrane was blocked for 30 minutes at room temperature in a 5% milk in TBST solution (Tris-buffer saline solution 1X, 0.05% Tween 20) followed by overnight incubation with the mouse anti-Vinculin V-9131 (Sigma) primary antibody in 5% milk in TBST solution. The membrane was then washed three times with TBST solution for 5 minutes each time and incubated for 1 hour at room temperature with HRP-Goat anti-Mouse (Invitrogen 626520) secondary antibody in 5% milk in TBST solution. Prior to revelation the membrane was once again washed three times with TBST solution for 5 minutes each time. Signal was developed using SuperSignal® West Femto Maximum Sensitivity Substrate Kit (Thermo Scientific) according to the manufacturer's instruction and a ChemiDoc™ MP Imaging system (BioRad).

**Statistical analysis.** Statistical analyses were performed in GraphPad Prism VII, R or Python. Data are represented with mean ± SEM unless otherwise stated. Data were pooled between independent replicates whenever the variability between statistical units within an experiment accounted for the total variability. Differences between the control FL and mutants were assessed with the non-parametric, Kruskal-Wallis and unpaired, two-tailed Dunn's tests. Differences between front and back cells were assessed with the non-parametric, two-tailed Mann-Whitney test. Multiple comparisons were accounted for by correcting p-values with the Benjamini-Krieger-Yekutieli (BKY) two-stage step-up FDR procedure after a positive-dependency check between comparisons by bootstrapping. Linear regressions by the least-squares method were tested against the null hypothesis slope=0 using an extra sum-of-squares F test. FRAP fits by the least-squares method were compared using an extra sum-of-squares F test with shared immobile fraction between FL and each mutant as the null hypothesis. Multiple comparisons were also accounted for by correcting p-values with the BKY procedure. Correlations were calculated using the Pearson correlation. Changes in single cell migration speed $\tilde{\Delta} v_{\text{s}}$, in difference between single cell and collective migration speeds $\tilde{\Delta} v_{\text{s-c}}$ and in ratio between wound healing rate and collective migration speed $\tilde{\Delta} v_{\text{c/d}}$ across vinculin constructs were compared to corresponding changes in vinculin tension $\tilde{\Delta} T_{\text{FA/AJ}}$, stability $\tilde{\Delta} S_{\text{FA/AJ}}$ and tension gradients $\tilde{\Delta} m_{\text{FA/AJ}}$ by re-scaling $v_{\text{s}}$, $v_{\text{s}} - v_{\text{c}}$, $v_{\text{c}}/v_{\text{d}}$, $1 - I_{\text{F, FA/AJ}}$, $F_{\text{im, FA/AJ}}$ and $-\Delta I_{\text{F,FA/AJ}}$ respectively, between their extreme values across vinculin constructs (e.g. $\tilde{\Delta} v_{\text{s}} = (v_{\text{s}} - \min(v_{\text{s}}))/(\max(v_{\text{s}}) - \min(v_{\text{s}}))$. Linear regression by the least-squares on $\tilde{\Delta} v_{\text{s}}$ or $\tilde{\Delta} v_{\text{s-c}} = w\tilde{\Delta} T_{\text{FA/AJ}} + (1-w)\tilde{\Delta} S_{\text{FA/AJ}}$ was used to determine the weight $w \in [0,1]$. Because $w = 0$ was the optimum in the second case, we then relaxed the linearity on $\tilde{\Delta} v_{\text{s-c}} = \tilde{\Delta} S_{\text{AJ}}$ with $\tilde{\Delta} v_{\text{s-c}} = (\tilde{\Delta} S_{\text{AJ}})^{\beta}$. Linear regression by the least-squares on $\tilde{\Delta} v_{\text{c/d}} = \tilde{\Delta} m_{\text{FA}} - \beta \tilde{\Delta} m_{\text{AJ}}$ was used to determine the factor $\beta$. MSD models were compared using the likelihood score $1/(\exp \Delta\text{AIC} + 1)$ where AIC is the Akaike Information Criterion. *: p<0.05, **: p<0.01, ***: p<0.001, ****: p<0.0001. No method was used to define sample size or for blinding.

**Table 1.** Most likely walk model for single cell migration

| Cells | Likelihood score of persistent random walk vs. pure or anomalous diffusion | $P$(s) | $D(\mu m^2/s)$ |
|---|---|---|---|
| WT | >0.9999 | $578 \pm 12$ | $0.088 \pm 0.0004$ |
| FL | >0.9999 | $813 \pm 25$ | $0.045 \pm 0.0004$ |
| TL | >0.9999 | $1059 \pm 13$ | $0.0277 \pm 0.0001$ |
| T12 | >0.9999 | $970 \pm 15$ | $0.1217 \pm 0.0006$ |
| YE | >0.9999 | $567 \pm 58$ | $0.08 \pm 0.0004$ |
| RPE-1 | >0.9999 | $3956 \pm 114$ | $0.3468 \pm 0.0006$ |

**Table 2.** Most likely walk model for collective cell migration

| Cells | Likelihood score of persistent random walk vs. pure or super-diffusion | Anomalous exponent $\alpha$ |
|---|---|---|
| WT | $< 10^{-4}$ | $1.73 \pm 0.003$ |
| FL | $< 10^{-4}$ | $1.699 \pm 0.008$ |
| TL | $< 0.4172$ | $1.52 \pm 0.01$ |
| T12 | $< 10^{-4}$ | $1.545 \pm 0.007$ |
| YE | $< 10^{-4}$ | $1.636 \pm 0.007$ |
| $\alpha-$catenin KD | $< 0.1155$ | $1.192 \pm 0.009$ |
| E-cadherin KO | $< 10^{-4}$ | $1.847 \pm 0.005$ |
| RPE-1 | $< 10^{-4}$ | $1.625 \pm 0.005$ |

**Table 3.** Most likely walk model for directed collective cell migration

| Cells | Likelihood score of persistent diffusion vs. pure or super-diffusion | Anomalous exponent |
|---|---|---|
| WT | $< 10^{-4}$ | $1.52 \pm 0.01$ |
| FL | $< 10^{-4}$ | $1.718 \pm 0.003$ |
| TL | $< 10^{-4}$ | $1.571 \pm 0.008$ |
| T12 | $< 10^{-4}$ | $1.687 \pm 0.006$ |
| YE | $< 10^{-4}$ | $1.55 \pm 0.01$ |
| $\alpha-$catenin KD | $< 3.10^{-4}$ | $1.199 \pm 0.008$ |
| E-cadherin KO | $< 10^{-4}$ | $1.846 \pm 0.004$ |

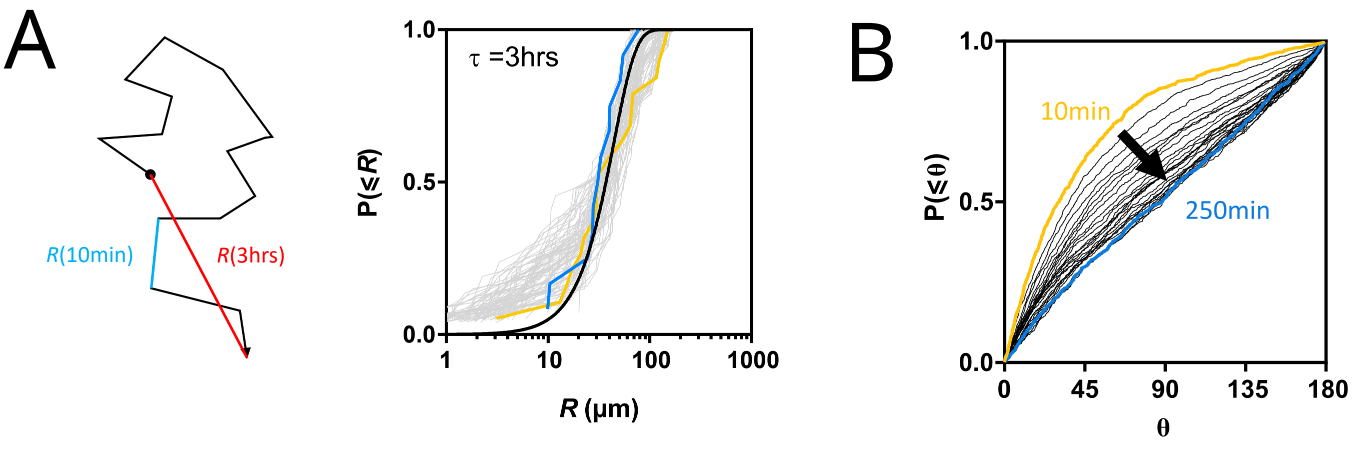


**Fig. S 1.** (A) Left: Schematic migration track and step length ($R(\Delta t)$, see Materials and Methods) at $\Delta t = 10$min (blue) and $\Delta t = 250$min (red). Right: cumulative distributions of the step length $R$ for $\Delta t = 3$hrs of single cells. One grey curve per frame pair along a movie. The yellow and blue curves correspond to the first and last frames, respectively. The black line is a global fit of all grey curves with a cumulative Rayleigh distribution function. (B) Cumulative distributions of the angular deviation ($\theta(\Delta t)$, see Materials and Methods) between $\Delta t = 10$min (yellow) and $\Delta t = 250$min (blue) of single cells. Data from 94 cell tracks of 2 independent experiments (same as in Fig. 1).

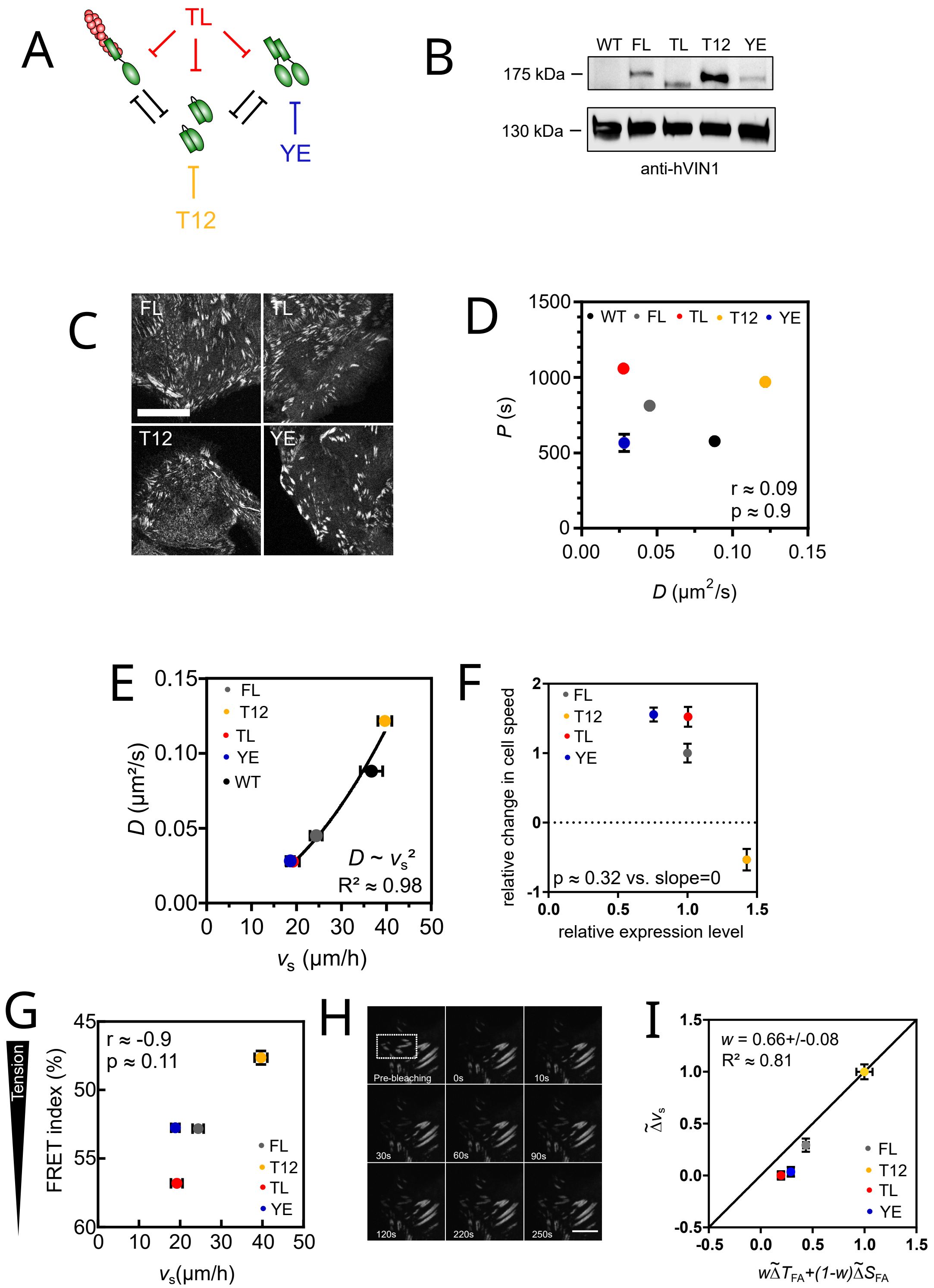


**Fig. S 2.** (A) Vinculin (green) and the different interactions (head-tail, tail-tail and actin-tail) and mutants (TL, T12 and YE) considered in this study. (B) Western Blot of the WT cells and cells expressing the FL, TL, T12, and YE vinculin TSMod constructs labelled for vinculin (WT at 130kDa, TSMod at 175kDa). (C) Direct fluorescence of the vinculin TSMod constucts in FAs. Scale bar=20$\mu$m. (D) Persistence time $P$ as a function of diffusion coefficient $D$ for WT, vinculin FL and mutant single cells. Pearsons correlation coefficient r and p-value. (E) Diffusion coefficient $D$ as a function of the apparent speed $v_s$ of WT, vinculin FL and mutant single cells, globally fit the Fürth equation with $\Delta t = 10$min. (F) Relative change in single cell apparent speed as a function of vinculin FL or mutant expression level. p-value of an extra sum-of-squares F test on a linear regression with slope=0 as the null hypothesis. The dashed line indicates the WT cell line as a reference. (G) FRET index as a function of apparent speed $v_s$ for vinculin FL and mutant single cells. Pearsons correlation coefficient r and p-value. (H) Direct fluorescence of vinculin TSMod in FAs during FRAP. White Box shows the FRAPed region. Scale bar=5$\mu$m. (I) Change in single cell speed $\tilde{\Delta} v_s$ as a function of weighted changes in tension $\tilde{\Delta} T_{FA}$ and $\tilde{\Delta} S_{FA}$, obtained from re-scaled means of $v_s$, $1 - I_{F,FA}$ and $F_{im,FA}$, respectively (data from Fig.2B-E). The weight $w$ maximizes the fit to the straight line of slope 1 through the origin. Data of (D-F) from 94 (WT), 125 (FL), 129 (TL), 193 (T12) and 110 (YE) cell tracks of 2 (WT) and 3 (all others) independent experiments (same as in Figs. 1 and 2A,B). Data of (G) as above for $v_s$ and from 81 (FL), 102 (TL), 52 (T12) and 84 (YE) cells of 4 (FL) and 3 (TL, T12, YE) independent experiments for FRET index (same as in Fig. 2B,D). Mean $\pm$ SEM.

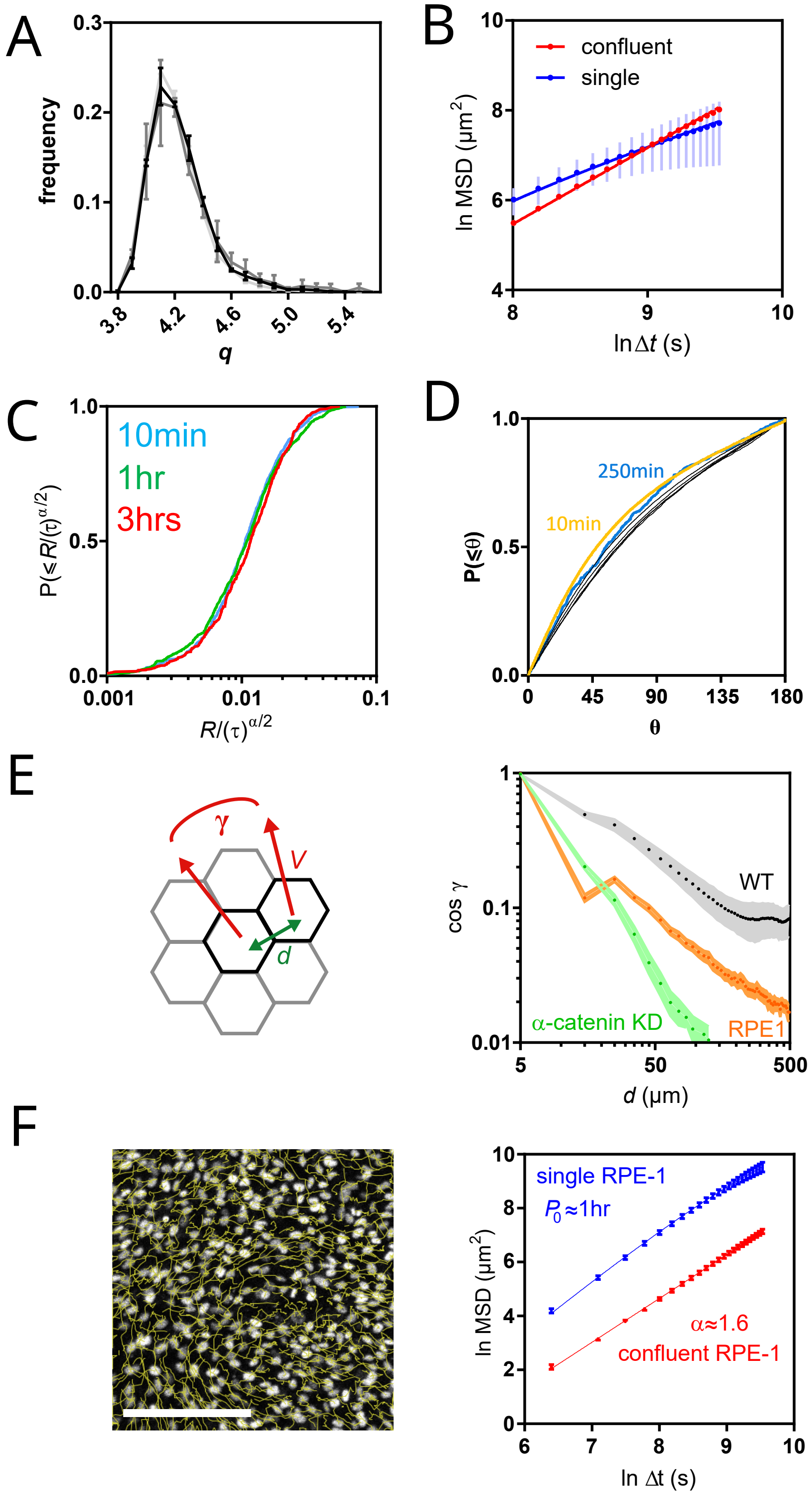


**Fig. S 3.** (A) Cell shape index index distribution of WT cells at confluence. The lines connect neighboring data points. (B) MSD($\Delta t$) of single (blue) and confluent (red) cells at long time scales fitted with a the Fürth model and a straight line in ln-ln scale, respectively. (C) Cumulative distributions of the normalized step length $R/(\tau)^{\alpha}$ where $\alpha$ is the anomalous exponent, for $\tau = 10$min, 1hr and 3hrs at confluence. One representative frame pair per $\tau$. (D) Cumulative distributions of the angular deviation $\theta(\Delta t)$, (see Materials and Methods) between $\Delta t = 10$min (yellow) and $\Delta t = 250$min (blue) of confluent cells. (E) Average coordination $< \cos\gamma >$ as a function of intercellular distance $d$ in WT and $\alpha$-catenin cells at confluence. (F) Left: fluorescence image of nucleus-stained RPE-1 cells at confluence and their migration tracks in yellow. Right: MSD($\Delta t$) of single (blue) and confluent (red) RPE-1 cells with best fitting models (colored lines) and their relevant parameters. Data from (A) 3 and (B-E) 11 (WT) and 4 ($\alpha$-catenin) FOV (500-1000 tracks each) of 3 independent experiments (same as in Fig. 3B-F). (F) Data from 11 FOV (500-1000 tracks each at confluence) of 2 (single) and 3 (confluent) independent experiments. Scale bar = 100μm. Mean $\pm$ SEM.

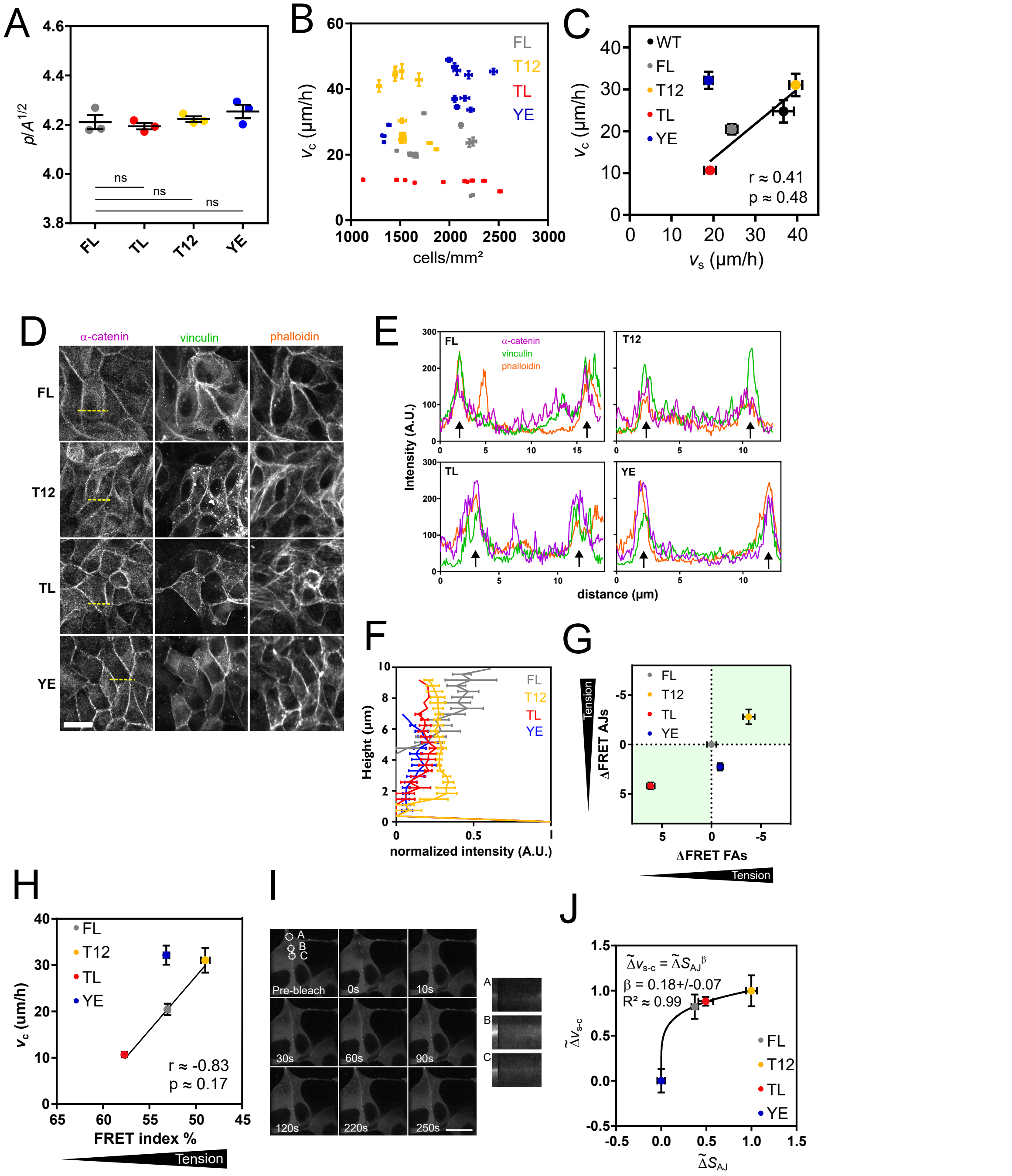


**Fig. S 4.** (A) Mean cell shape index as a fucntion of vinculin mutant. (B) Apparent collective cell speed ($v_c$) as a fucntion of cell density per FOV. (C) Apparent collective cell speed ($v_c$) as a function of apparent single cell speed ($v_s$) for WT cells and cells expressing the vinculin FL and mutant constructs. Pearson coefficient r and p-value for all conditions. The black line is a linear fit excluding YE. (D) Maximum intensity projections of immunofluorescence stainings of $\alpha$-catenin and actin (phalloidin) on FL, T12, TL and YE vinculin cell lines. Dashed lines indicate location of intensity profiles shown in (E). (E) Typical intensity profiles of immunofluorescence staining of $\alpha$-catenin and actin together with each vinculin construct along dashes lines shown in (D) (arrows indicate cell-cell contacts). (F) Lateral recruitment of vinculin constructs as a function of height along intercellular contacts normalized to FAs in live cells. Colored lines connect neighboring data points. (G) FRET index of the vinculin mutants relative to vinculin FL at AJs vs. FAs. (H) Apparent collective cell speed ($v_c$) as a function of FRET index of vinculin constructs. Pearson coefficient $R^2$ and p-value for all conditions. The black line is a linear fit without YE. (I) Left: direct fluorescence of vinculin TSMod in AJs during FRAP. Right: kymographs of the circled, FRAPed regions A, B, C on the left. (J) Change in the difference between single and collective cell migration speeds $\tilde{\Delta} v_{\text{s-c}}$ as a function of the change in vinculin stability in AJs $\tilde{\Delta} S_{\text{AJ}}$, obtained from re-scaled means of $v_s - v_c$ and $F_{\text{im,AJ}}$, respectively (data from Fig. S4C, 4F). The line is the fit with exponent $\beta$ on $\tilde{\Delta} F_{\text{im,AJ}}$. Data of (A) from 3 independent experiments for each mutant (500-1000 cells each). Data of (B,C) from 94 (WT), 125 (FL), 129 (TL), 193 (T12) and 110 (YE) cell tracks of 2 (WT) and 3 (all others) independent experiments for $v_s$) (same as in Figs. 1 and 2A,B) and from 11 (WT), 12 (FL), 11 (TL, T12) and 14 (YE) FOV (500-1000 tracks each) of 3 independent experiments for $v_c$) (same as in Figs. 3B-D and 4A,B). Data of (F) from 5(FL), 6(TL), 7(T12), 11(YE) cells. Data of (G) from 169 (FL), 30 (TL) and 64 (T12, YE) contacts of 5 (FL) and 3 (TL, T12, YE) independent experiments for AJs, and from 45 (FL), 35 (TL), 104 (T12) and 67 (YE) cells of 4 (FL) and 3 (TL, T12, YE) independent experiments for FAs (same as in Fig. 4D,E). Data of (H) as in (C) for $v_c$ and in (G, FAs) for FRET index. Mean $\pm$ SEM. Scale bar=20$\mu$m.

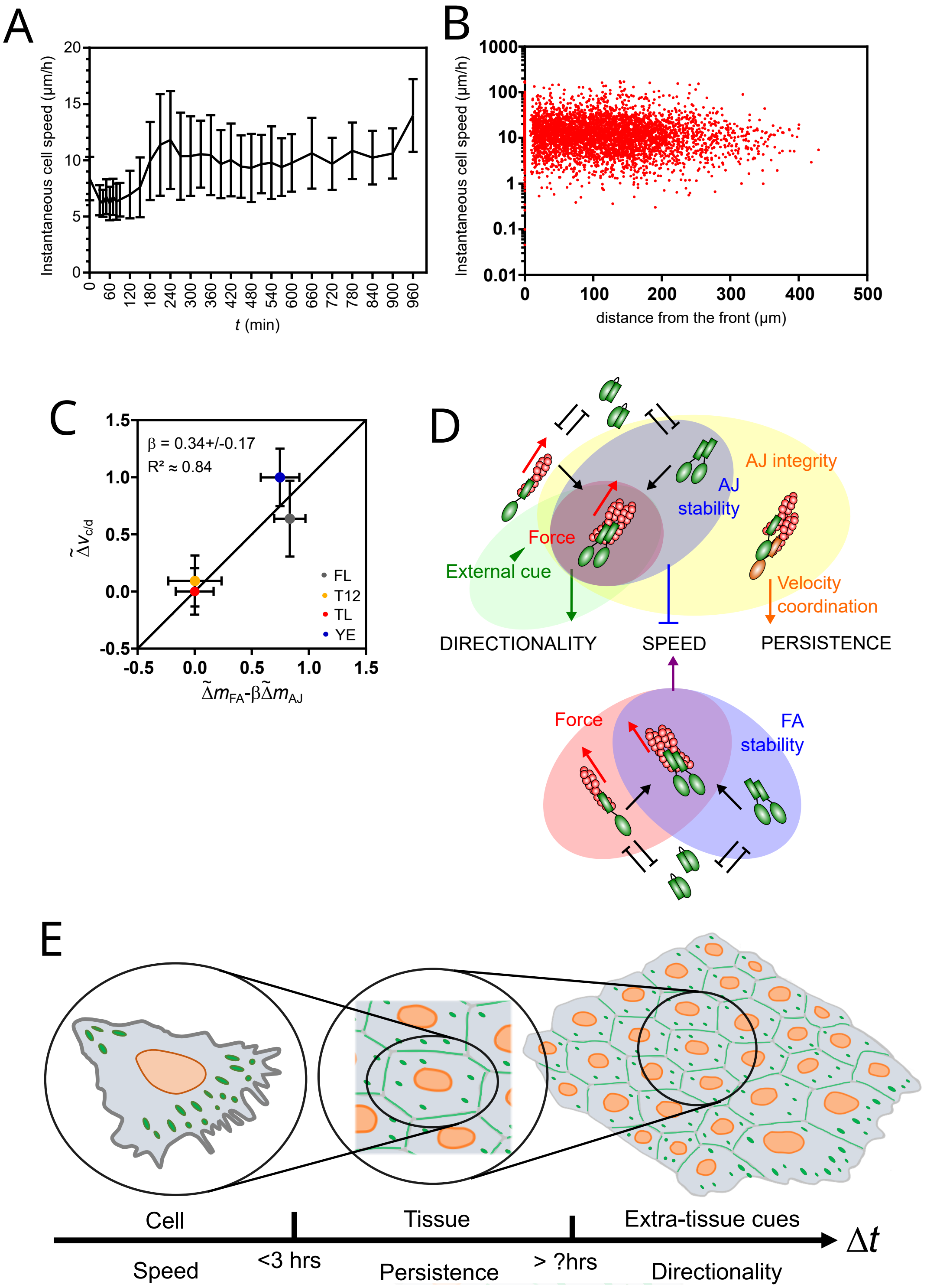


**Fig. S 5.** (A) Apparent cell speed as a function of time in a wound healing assay. The black line connects neighboring data points. (B) Apparent cell speed as a function of the distance to the front at 6 hours after wounding. (C) Change in the ratio of collective cell migration speed to wound healing rate $\tilde{\Delta} v_{c/d}$ as a function of the difference of mechanosensitivity to the wound between FAs $\tilde{\Delta} m_{FA}$ and AJs $\tilde{\Delta} m_{AJ}$, obtained from re-scaled means of $v_c/v_d$, $-\Delta I_{F,FA}$ and $-\Delta I_{F,AJ}$, respectively (data from Fig.6A,C,E). The factor $\beta$ maximizes the fit to the straight line of slope 1 through the origin. (D) Complete Boolean model for the control of cell speed persistence and directionality by the stability and force transmission of adhesions, from Fig. 2F, 4G, 6F). (E) Working model for the control of cell spatial exploration efficiency as a function of time scales (see text for details). Data of (A) from 3 FOV (500-1000 tracks each) of 3 independent experiments. Mean ± SEM. Data of (B) from 5 FOV (about 1000 cells each) of 5 independent experiments.
